# β-Carotene oxidation products - function and safety

**DOI:** 10.1101/2021.03.18.436000

**Authors:** Graham W. Burton, Trevor J. Mogg, William W. Riley, James G. Nickerson

## Abstract

β-Carotene oxidation products have newly discovered bioactivity in plants and animals. Synthetic fully oxidized β-carotene (OxBC) has application in supporting livestock health, with potential human applications. The safety of synthetic OxBC has been evaluated. An Ames test showed weak-to-moderate mutagenicity in only one cell line at high concentrations. A mouse micronucleus assay established a non-toxic dose of 1800 mg/kg body weight, and no bone marrow micronuclei were induced. Plant sources of β-carotene inevitably contain varying levels of natural OxBC. Vegetable powders and dried forages can be especially rich. Intakes of natural OxBC for humans and livestock alike have been estimated. The exposure range for humans (1-22 mg/serving) is comparable to the safe intake of β-carotene (<15 mg/d). In livestock, OxBC in alfalfa can contribute ~550-850 mg/head/d for dairy cattle but in forage-deficient poultry feed much less (~1 ppm). Livestock intake of supplemental synthetic OxBC is comparable to OxBC potentially available from traditional plant sources. Human intake of synthetic OxBC in meat from livestock fed OxBC is similar to a single serving of food made with carrot powder. It is concluded that consumption of synthetic OxBC at levels comparable to natural OxBC is safe for humans and animals.

## 1. Introduction

The common underlying polyene electronic structure that gives rise to the strikingly intense colorations of β-carotene and the other carotenoids makes the compounds quite susceptible to oxidation. In Nature enzymatic and non-enzymatic carotenoid oxidations generate products with diverse properties that can include physiological and sensory activities. Important examples of enzymatically generated products include vitamin A in animals and the abscisic acid and strigolactone hormones in plants. Non-enzymatic oxidation of carotenoids in plants contributes a cornucopia of apocarotenoid products, many with sensory properties finding applications as fragrances and flavors (Winterhalter and Rouseff, 2001).

There is a growing body of evidence that β-carotene oxidation products other than vitamin A or plant hormones can contribute significant biological functions in plant and animal physiology. For example, in plants the apocarotenoid ß-cyclocitral has been identified as playing an important protective signaling role when generated at very low concentrations via photochemical quenching by β-carotene under excessive sunlight exposure conditions (D’Alessandro et al., 2018). This action exploits the highly reactive β-carotene as an oxidation sensor, indirectly triggering gene expression that activates defense mechanisms to prevent harmful cellular oxidation.

In humans there has been concern that some of β-carotene’s larger, retinoid-like β-apocarotenoid oxidation products are potentially toxic. These compounds became implicated in the unexpected negative outcomes of the large-scale β-Carotene and Retinol Efficacy (CARET) Trial and the α-Tocopherol, β-Carotene Cancer Prevention (ATBC) Study involving daily supplementation with 20-30 mg β-carotene (ATBC Cancer Prevention Study Group, 1994; Goodman et al., 2004; Omenn et al., 1994; Omenn et al., 1996a; Omenn et al., 1996b; Virtamo et al., 2014). The high-dose β-carotene supplements increased the risk of lung cancer among smokers and asbestos workers.

These findings stand in stark contrast to those of observational epidemiologic studies that have consistently shown individuals who eat more fruits and vegetables, rich in carotenoids, and people with higher serum β-carotene levels have a lower risk of cancer, particularly lung cancer (Mayne, 1996). Earlier, it had been hypothesized that β-carotene itself could be contributing a preventive antioxidant effect independent of its vitamin A activity (Peto et al., 1981).

In a third human intervention trial, the Physicians’ Health Study, conducted mainly among non-smokers, no effect, however, was found on lung cancer risk in either smokers or non-smokers given 50 mg of β-carotene every other day (Hennekens et al., 1996). It has been noted there were far fewer smokers in this trial and that the serum and tissue β-carotene levels were much higher in the ATBC and CARET trials (Russell, 2004).

In attempting to better understand the confounding trial results, focus has been directed to applying a variety of *in vitro* toxicity assays to β-carotene oxidation products generated by simulating presumed *in vivo* oxidative conditions (Alija et al., 2010; Alija et al., 2005, 2006; Alija et al., 2004; Eroglu et al., 2012; Hurst et al., 2004; Hurst et al., 2005; Kalariya et al., 2008, 2009, 2011; Marques et al., 2004; Salerno et al., 2007; Salgo et al., 1999; Siems et al., 2002; Siems et al., 2000; Siems et al., 1999; Sommerburg et al., 2003; Yeh and Hu, 2003; Yeh and Wu, 2006). A majority of the studies used partially oxidized β-carotene mixtures obtained by reaction with hypochlorous acid. However, the possibility exists that the reaction products included potentially genotoxic chlorinated compounds. The other *in vitro* studies used partially oxidized β-carotene prepared according to the air-oxidation method of Handelman et al. (Handelman et al., 1991). Notably, several longer chain β-apocarotenoid cleavage products were identified as possible toxicity candidates in both types of simulated oxidized β-carotene product mixtures (Alija et al., 2005; Alija et al., 2004; Eroglu et al., 2012; Kalariya et al., 2009; Marques et al., 2004; Yeh and Wu, 2006). The physiological relevance of these *in vitro* studies has been questioned in a review in 2012 of the safety of β-carotene by an EFSA Panel (European Food Safety Authority, 2012a).

The clearest insight into the origins of the conflicting β-carotene clinical trial results has come from studies using a ferret smoking inhalation model, which reproduced the three human intervention trial results (Russell, 2004; Wang et al., 1999). With smoke exposure high-dose β-carotene led to pre-cancerous, squamous metaplasia lesions in the ferret lung, whereas low dose β-carotene provided mild protection against squamous metaplasia. The effects of β-carotene on lung cancer therefore appear to be related to the β-carotene dose.

Cigarette smoke exposure decreased the levels of β-carotene in both plasma and lung in ferrets given the β-carotene supplement and in the lung in the unsupplemented group (Wang et al., 1999). *In vitro* incubations of β-carotene with post-nuclear fractions of lung tissue from ferrets exposed or not exposed to smoke showed enhanced formation of β-apocarotenoid oxidative metabolites associated with the decrease in β-carotene. The levels of β-apo14’, β-apo12’, β-apo10’, and β-apo8’ carotenals were threefold higher in lung extracts of smoke-exposed ferrets than in lung extracts of non-exposed ferrets. The free radical rich, oxidative atmosphere in the lungs of cigarette smoke exposed-ferrets affected β-carotene metabolism, leading to increased levels of β-apocarotenoid metabolites that are structurally similar to retinoids and that have been found to interfere with the metabolism of retinoic acid and retinoid signaling.

In responding to a related concern in connection with the possibility of high levels of β-carotene in β-carotene-biofortified plant crops, e.g., Golden Rice, a comprehensive study of β-carotene degradation in a wide range of conventional and biofortified post-harvest crop food items found that the quantity of apocarotenoids generated by oxidative degradation during storage was two orders of magnitude *less* than the loss of β-carotene (Schaub et al., 2017). Total levels of apocarotenoids remained very low (ng/g) compared to changes in β-carotene levels (μg/g). Significantly, a β-carotene-oxygen copolymer was found to account for the discrepancy in the degradation product yield, corroborating our earlier finding that the copolymer is the main non-enzymatic oxidative species in dried plant products (Burton et al., 2016). It was concluded that the very low levels of apocarotenoids associated with the dietary intake of conventional or β-carotene-biofortified plant items do not pose a human safety risk.

In livestock animals it had been thought for some time that β-carotene is a source of additional biological activity unrelated to its provitamin A function. However, in 2012 in a review of the safety and efficacy of β-carotene as a feed additive for all animal species, an EFSA Panel concluded that non-vitamin A effects, for example on reproduction and immunity, had not yet been sufficiently demonstrated (European Food Safety Authority, 2012b).

Subsequently, a comprehensive, detailed approach to the spontaneous oxidation of β-carotene has brought a degree of clarity and unity to the disparate findings associated with β-carotene’s apparent non-vitamin A activities (Burton et al., 2014; Mogg and Burton, 2020). Focusing on the complete mixture of the many oxidation products generated by the full spontaneous oxidation of β-carotene revealed the existence of compounds capable of non-vitamin A antiproliferative and anti-tumor activity (Burton et al., 1995) and immunological activity (Duquette et al., 2014; Johnston et al., 2014).

Full oxidation of pure β-carotene in air reproducibly generates a mixture of compounds referred to as synthetic OxBC that is comprised of a small quantity of a complex mixture of many apocarotenoid breakdown products together with a major quantity of a previously unknown β-carotene-oxygen copolymer product (Burton et al., 2014). This same oxidation reaction occurs in carotenoid-containing crop plant products during storage or drying (Burton et al., 2016; Schaub et al., 2017) and has been accompanied by the isolation of carotenoid-oxygen copolymer compounds in several examples. It is noteworthy that synthetic OxBC does not contain the potentially toxic, long-chain apocarotenoids (Burton et al., 2014). OxBC’s cleavage products are all small chain apocarotenoids containing 8 to 18 carbon atoms, less than half of β-carotene’s 40 carbons, with only a few present at the 1% level (by weight) and the rest at much less than 1%. The more reactive, long-chain apocarotenoid products (≥C_20_) initially formed are ultimately consumed in the full oxidation reaction. Thirteen of the apocarotenoids available in OxBC have been designated as Generally Recognized As Safe (GRAS) human flavor agents (U.S. Food & Drug Administration).

Synthetic OxBC has proven to be a useful tool for screening and probing the biological activity of β-carotene oxidation products. Lacking β-carotene, vitamin A and vitamin A activity, the demonstration of OxBC biological activity in both *in vitro* and *in vivo* mammalian assays has provided convincing evidence that the source of non-vitamin A activity exists within the oxidation products themselves (Johnston et al., 2014).

From a practical viewpoint, livestock trials with low, parts-per-million (ppm) level supplementation of synthetic OxBC in feed have shown performance benefits over and above the benefits of feeds containing regular vitamin and mineral premixes (Chen et al., 2020; Kang et al., 2018; McDougall, 2020). When the introduction of synthetic vitamin A replaced provitamin A forages, β-carotene and its oxidation products largely disappeared from livestock feeds.

Supplementing feeds with synthetic OxBC can restore the previously unrecognized benefits of β-carotene oxidation products.

Several of OxBC’s small apocarotenoids are reactive electrophiles containing α,β-unsaturated carbonyl groups, e.g., β-cyclocitral, or other reactive carbonyls, e.g., methyl glyoxal (Mogg and Burton, 2020). Very recently Koschmieder, Welsch and coworkers demonstrated that plants modified to express excess β-carotene exhibit metabolism of the apocarotenoid products utilizing defense mechanisms against reactive carbonyl species and xenobiotics (Koschmieder et al., 2020). Earlier, Schaub and coworkers, employing a non-green *Arabidopsis callus* system, identified oxidative β-carotene degradation products, among them β-apocarotenals of different chain lengths and several putative end-product dialdehydes, methylglyoxal and glyoxal (Schaub et al., 2018). These latter compounds have independently been shown to be released from the OxBC copolymer (Mogg and Burton, 2020).

Often, compounds with a botanical origin such as those in OxBC have a documented history of exposure and use. However, the significance of naturally occurring OxBC and the copolymer have only recently been recognized (Burton et al., 2016). Although long-time human and livestock exposure to naturally occurring OxBC consumed in plant-derived food and feed components implies a degree of safety, the presence of the previously unrecognized copolymer together with small amounts of potentially reactive compounds requires a more critical assessment of the safety of the substance.

In this paper the safety of β-carotene oxidation products, in the form of synthetic OxBC’s mixture of small apocarotenoids and the β-carotene copolymer compound, is evaluated directly in genotoxicity assays and less directly through a literature survey to estimate dietary exposure to natural OxBC from various foods and feeds consumed by humans and animals alike.

## 2. Materials and methods

### 2.1. Genotoxicity tests

All genotoxic studies were performed by Charles River Laboratories, Tranent, Edinburgh, UK. The studies were conducted in compliance with OECD, European Commission and U.S. Environmental Protection Agency (EPA) guidelines, and in accordance with the OECD Principles of Good Laboratory Practice.

The mouse studies were conducted in accordance with the UK Home Office Legislation. The care and use of mice conformed with the U.K. Animals (Scientific Procedures) Act, 1986 and associated guidelines, Home Office Licence Number PPL 60/3517, Toxicology of Medical and Veterinary Materials, Procedure 3 (toxicity study) and Procedure 4 (micronucleus test).

#### 2.1.1. Test article

OxBC was prepared commercially by Allied Biotech, Taipei, by oxidation of pure synthetic β-carotene in a solvent under an atmosphere of pure oxygen. The properties have been described elsewhere (Burton et al., 2014). The test item, OxBC, was stored as a solution in dimethyl sulfoxide (DMSO), unless noted otherwise, at −20°C when not in use

#### 2.1.2. Bacterial reverse mutation assay (Ames test)

OxBC was tested for mutagenic activity in *Salmonella typhimurium* strains TA 1535, TA 1537, TA 98 and TA 100 and in *Escherichia coli* WP2*uvrA*. Dimethyl sulfoxide (DMSO) was used as the solvent and vehicle for OxBC with test formulations being prepared immediately prior to dosing (within 1 h). Mutagenic activity was assessed with the bacterial reverse mutation assay (U.S. Food and Drug Administration, 2007). An S9 mixture (S9), a cytosolic homogenate prepared from the livers of Aroclor 1254-treated rats, along with cofactors necessary for enzymatic activity, provided an exogenous metabolic activation system (McGregor et al., 1988).

Two experiments were conducted in both the absence and the presence of S9. OxBC was dosed at concentrations spaced at half-log intervals in the first mutation assay and at concentrations spaced at halving intervals in the second mutation assay. A toxicity test was performed to establish the concentration range to be used in the first mutation test, covering 17, 50, 167, 500, 1667 and 5000 μg per plate. The positive controls were 2-aminoanthracene, sodium azide, N-ethyl-N-nitro-N-nitrosoguanidine, 2-nitrofluorene and 9-aminoacridine.

##### Interpretation of Mutagenicity

For *S. typhimurium* strains TA 1535 and TA 1537 and for *E. coli* WP2*uvrA*, at least a 3-fold increase over the mean concurrent vehicle control value was required before mutagenic activity was suspected. For *S. typhimurium* strains TA 98 and TA 100, a 2-fold increase over the control value was considered indicative of a mutagenic effect. A concentration-related response was also required for identification of a mutagenic effect.

The experimental procedure is described in more detail in Supplementary Material S1.

#### 2.1.3. Chromosomal aberration assay

This assay tested the ability of OxBC to induce chromosomal aberrations in cultured Chinese hamster ovary cells. The study complied with OECD and ICH Guidelines and with the European Commission Annex V, Test Method B10. The experimental procedure is described in detail in Supplementary Material S1.

#### 2.1.4. Mouse micronucleus assay

OxBC was evaluated for *in vivo* clastogenic activity and/or disruption of the mitotic apparatus by detecting micronuclei in polychromatic erythrocytes in CD-1 mouse bone marrow. Experimental procedures complied with OECD, ICH and EC Guidelines, US EPA Pesticide Assessment Guidelines, and the Japanese Guidelines on Genotoxicity Testing and recommendations published by the US EPA Gene-Tox Program and the Japanese Collaborative Study Group for Micronucleus Testing (Environmental Mutagen Society, 1990; Mavournin et al., 1990). The vehicle control was a solution of N-methyl-2-pyrrolidone, polyethylene glycol 400 and propylene glycol prepared in the ratio of 1:2:2, respectively. The experimental procedure is described in detail in Supplementary Material S1.

### 2.2. OxBC exposure from feeds, foods and supplements

Use of geronic acid (GA) as a marker of OxBC content in a variety of vegetable powders (Burton et al., 2016) has allowed estimation of OxBC levels in a variety of human foods prepared using these powders. Similarly, knowledge of the level of OxBC in dried forages and other livestock feed items has permitted an estimation of the level of exposure of livestock to OxBC. A literature survey was undertaken to gauge the extent of OxBC exposure in humans and livestock from the use of vegetable powders in foods and forages in livestock feeds, respectively.

## 3. Results

### 3.1 Bacterial reverse mutation assay (Ames Test)

OxBC showed weak to moderate activity in only the *Salmonella typhimurium* TA 100 strain when tested to the predetermined maximum concentration of 5000 μg per plate with metabolic activation and at concentrations extending into the toxic range without metabolic activation (Table 1). The mutagenicity threshold of a 2-fold excess of revertants over control was reached at 500 μg per plate without metabolic activation and between 500 and 1667 μg per plate with metabolic activation. No mutagenic activity was detected in the three other *Salmonella* strains, TA 1535, TA 1537, TA 98, nor in the *Escherichia coli* WP2*uvrA* strain.

**Table 1.**
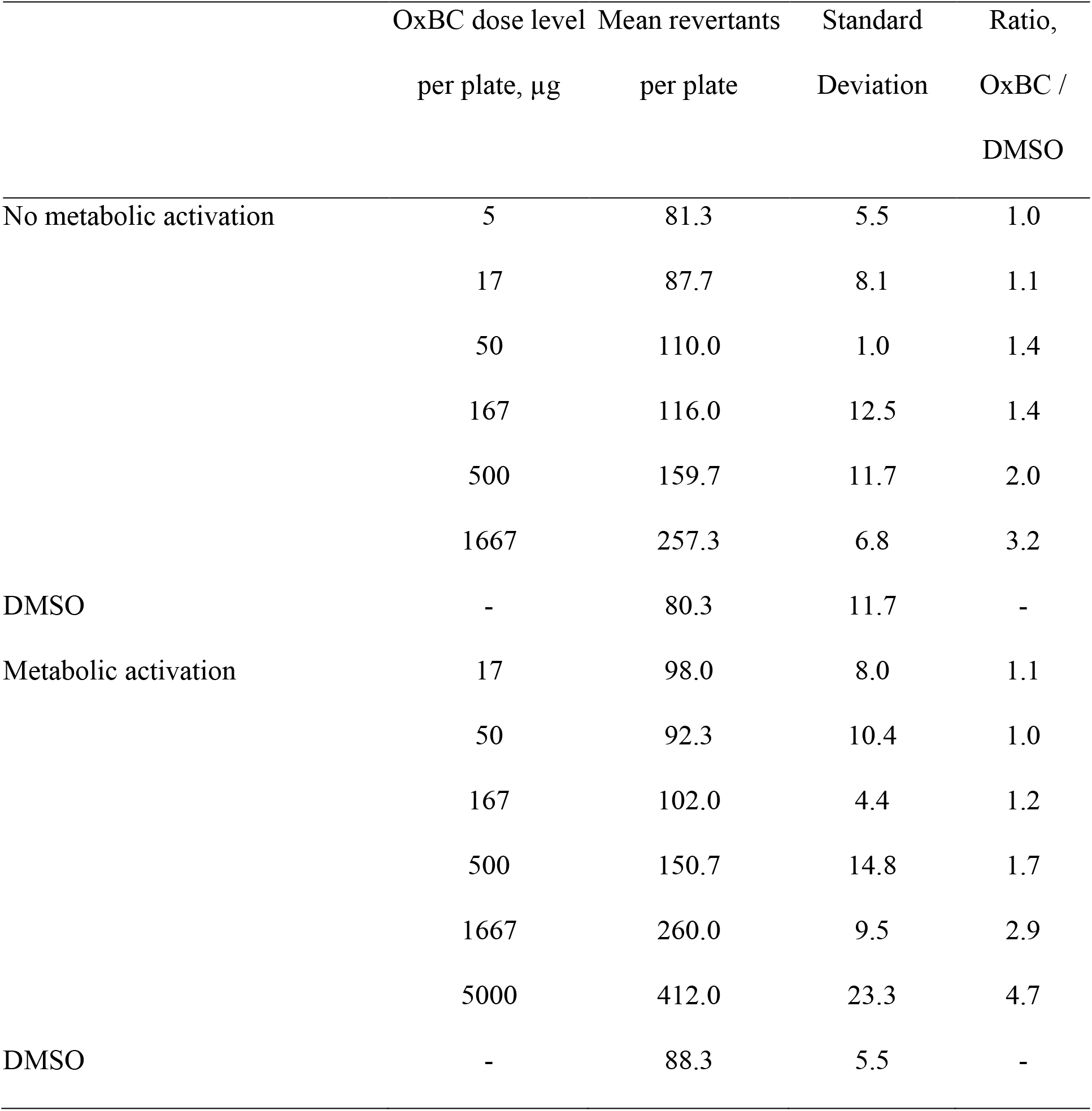
Mutation results from OxBC dosing of *Salmonella typhimurium* strain TA 100 without and with metabolic activation with S9.

Toxicity to the bacteria was generally observed as a thinning of the background lawn of microcolonies at the higher doses. In the first mutation assay, toxicity to the bacteria was observed as a thinning of the background lawn of microcolonies at 1667 μg per plate in the absence of S9, and at 5000 μg per plate in the presence of S9, in strains TA 1535, TA 1537 and TA 100. In the second mutation assay, toxicity to the bacteria was observed at 2500 μg per plate in the absence of S9, and at 5000 μg per plate in the presence of S9, for all *S. typhimurium* strains. Precipitation of OxBC occurred at the highest concentration tested, 5000 μg per plate. The vehicle control values for the accepted tests were within the normal historical ranges from the laboratory and reported in the literature for these strains of *S. typhimurium* and *E. coli* (Ames et al., 1975; Gatehouse et al., 1994). The positive control values were within normal historical ranges for this laboratory for each strain and activation condition.

### 3.2 Chromosomal aberration assay

OxBC was clastogenic when tested with Chinese hamster ovary cells *in vitro* at concentrations deemed toxic to the cells, with structural chromosomal aberrations induced in both the presence and absence of S9. In the presence of S9, toxicity was noted in the treated cultures ranging from 78-5000 μg/mL, with reduced cell counts (below 50% of the vehicle control) in those cultures treated with 156-5000 μg/mL. In the absence of S9 (6 and 22 h treatments), toxicity was noted in the treated cultures ranging from 78-5000 μg/mL, with reduced cell counts at these concentrations. There was an indication of toxicity from the culture observation at 39 μg/mL at the longer treatment time. In the aberration test, in the presence of S9, toxicity was noted in the cultures treated with 40-100 μg/mL. Reduced cell counts, with no metaphase cells for assessment, were observed in cultures treated with 60-100 μg/mL. Concentrations of 40 and 50 μg/mL were deemed cytotoxic from slide observations, with mean cell counts of 63% and 62%, respectively, compared to vehicle controls. In the absence of S9, toxicity was noted in cultures treated with 20-90 μg/mL. Reduced cell counts were noted in cultures treated with 30-90 μg/mL, with too sparse or no metaphase cells for assessment in cultures treated with 50-90 μg/mL. Cultures treated with 20 μg/mL had a mean cell count of 58% compared to vehicle controls, and this concentration was deemed toxic to cells from slide observations.

### 3.3. Mouse micronucleus assay

OxBC did not induce micronuclei in bone marrow cells when tested to the maximum tolerated dose of 1800 mg/kg/day in male CD-1 mice using a 0 h + 24 h oral dosing and a 48-h sampling regimen. In a preliminary study to determine the maximum tolerable dose of OxBC, five groups of male and female CD-1 mice received oral doses of OxBC, ranging from 1600 to 2000 mg/kg/day. The following pattern of animal deaths occurred: Group 1 (1M + 1F) - 2000 mg/kg/day, 0 deaths; subsequently, Groups 2-4 (3M + 3F) - 2000 mg/kg/day, 2 deaths; Group 5 (3M + 3F) - 1600 mg/kg/day, 0 deaths.

The accompanying clinical signs were hunched posture, subdued behaviour, piloerection, staggering, laboured breathing, tremors, unwillingness to move, pale extremities and discolouration of urine. Based on these toxicity investigations, the maximum tolerated dose of OxBC was judged to be in the region of 1800 mg/kg/day. There was no evidence of a significant difference in toxicity between male and female mice, therefore only males were used in the micronucleus test.

In the micronucleus test, three groups of male CD-1 mice were dosed orally at 0 h + 24 h with OxBC at doses of 450, 900 and 1800 mg/kg/day. Concurrently, vehicle and positive control groups of mice were similarly dosed orally at 0 h +24 h and sampled at 48 h in parallel with the OxBC-treated mice. Clinical signs of hunched and subdued behaviour were observed. One animal was found dead before being dosed, but no animal deaths occurred during the test. The numbers of micro-nucleated bone marrow polychromatic erythrocytes (MN-PCE) in mice dosed with the vehicle at 10 mL/kg averaged 0.02%. In the untreated control group, the MN-PCE frequency averaged 0.02%. These MN-PCE frequencies conformed to the established in-house control range for vehicle treated mice of the CD-1 strain (0.00-0.23% per 5 mice). Exposure of mice to the positive control agent, 50 mg cyclophosphamide/kg, induced large increases in bone marrow micronuclei. The mean MN-PCE frequency for the mice was 1.26%. An evident increase in the number of micro-nucleated normo-chromatic erythrocytes (MN-NCE) was also observed. Bone marrow toxicity accompanied these findings, as shown by a slight suppression of the PCE/NCE ratios. There was no indication that OxBC induced bone marrow micronuclei in the treated mice. The highest MN-PCE frequency recorded for OxBC was in the high dose group, where an incidence of 0.04% was observed. As there was no indication of bone marrow toxicity in any of the OxBC dose groups it was concluded OxBC tested to the maximum tolerated dose did not induce micronuclei in bone marrow cells.

### 3.4. OxBC exposure from feeds, foods and supplements

#### 3.4.1. Livestock feeding trials

No adverse events have been reported in 31 feeding trials conducted under commercial conditions in various countries. Eleven feeding trials have been conducted in poultry involving 38,682 birds using mostly 2-4 ppm but also 5-30 ppm OxBC, 13 feeding trials involving 2,672 pigs using mostly 2-8 ppm but also 30-100 ppm OxBC, and seven feeding trials involving 345 dairy cows dosed at 0.25-0.35 g OxBC/head/day.

#### 3.4.2. Companion animals

Avivagen Inc., as a member of the United States National Animal Supplement Council (NASC), complies with the NASC set of standards, including Current Good Manufacturing Practices (cGMP), maintaining an Adverse Event Reporting System, and meeting labeling and claims guidelines set up in cooperation with the FDA. More than 1 million doses of OxBC formulations for companion animals, mostly dogs but also cats, have been provided over the 2010-2020 period. Only six adverse events have been reported, and none of these were serious.

#### 3.4.3. Exposure from dietary sources and supplements

Use of the GA marker compound has established the existence and extent of β-carotene oxidation, thereby enabling estimates of the levels of OxBC in a broad array of β-carotene-rich, plant-based food items (Burton et al., 2016). A model study of the production of carrot powder from carrot purée clearly shows the inverse relationship between the release of GA and the disappearance of β-carotene (Supplementary Material S2, Table 1 and Fig. 1).

Table 1 in Supplementary Material S3 provides updated values of estimated OxBC levels in a variety of plant products relative to previously reported estimates ((Burton et al., 2016). The updated values reflect a lower content of GA (~1%*vs*. ~ 2%) in the reference sample of OxBC prepared with air instead of pure oxygen to more realistically reflect the conditions under which natural OxBC forms.

The richest source of OxBC is carrot powder. As shown for the two commercial examples in Supplementary Material S3, Table 1, the estimated amounts of OxBC are comparable to the amounts of β-carotene copolymer compound isolated, which represents ~80% of OxBC by weight. The carrot powder sample with the highest OxBC content, approaching mg/g, was light brown and contained no measurable amount of β-carotene. Carotene copolymer isolated from carrot powder has been shown to be chemically very similar to the copolymer isolated from synthetic OxBC (Burton et al., 2016).

Similarly, mixed carotenoid copolymer compounds have been isolated from powders of tomato, rosehip, sun-cured alfalfa, dulse seaweed, wheatgrass and paprika. The total amounts of isolated carotenoid copolymers were substantially larger than the corresponding estimated levels of OxBC (Burton et al., 2016) (Supplementary Material S3, Table 1). Pure lycopene, lutein and canthaxanthin, as examples of other common carotenoids, very readily oxidize in a similar manner, forming carotenoid oxygen-copolymer compounds as the main product (Burton et al., 2016). These compounds show strong chemical similarity to the β-carotene copolymer.

The amounts of isolated carotenoid copolymer compounds in excess of OxBC copolymer reflect the relative abundances of the other carotenoids, especially lycopene and lutein, in the vegetable powders, as well as their relative tendencies to oxidize. In tomato powder, the amount of carotenoid copolymer is more than three times that in carrot powder, and it is estimated to be 90-fold higher than that of the α- + β-carotene copolymers also present (Supplementary Material S3, Table 1), even though lycopene is just approximately five times more abundant than the α- and β-carotenes together in tomato (United States Department of Agriculture, 2020). This observation is explained by the greater relative reactivity of lycopene toward oxidation and the even greater extent to which lycopene-oxygen copolymers are formed (Burton et al., 2016).

The information on the levels of OxBC in the vegetable powders listed in Supplementary Material S3, Table 1 has permitted an estimate of the exposure to OxBC from a variety of foods in which these powders are ingredients. The results of a literature survey presented in Supplementary Material S3 indicate that exposure of humans and animals to dried vegetable products, particularly carrot and tomato, has occurred over a very long time, spanning many centuries, and that currently there is widespread and extensive use of carrot and tomato powders in the food industry. Humans and livestock animals have had prolonged dietary exposure to significant levels of oxidized carotenoids, including OxBC, and their associated copolymers, especially since the late 1800s - early 1900s.

#### 3.4.4. Exposure to natural OxBC in humans

Dried vegetable ingredients are used extensively in a variety of foods by food manufacturers to prepare numerous products. For example, carrot, tomato and sweet potato powders are often used as ingredients to prepare baby foods, including instant meals, teething biscuits and snack puffs. Several manufacturers provide recipes on their websites that use significant quantities of carrot, sweet potato and tomato powder ingredients. Tables 2 and 3 in Supplementary Material S3 give the estimated amounts of OxBC and total carotenoid copolymers for a variety of common foods that contain significant amounts of dried vegetable or fruit ingredients. Baby food, baked goods, soups, stews and casseroles prepared with carrot powder are estimated to contain as much as 4-22 mg OxBC per single serving. A recipe for a drink or smoothie using carrot powder is estimated to contain approximately 4-7 mg of OxBC.

Carrot fiber, a specialized type of carrot powder, is an FDA-approved GRAS food additive (Bolthouse Farms, 2002) that is used extensively in the food industry. An off-white powder derived from the remains of fresh carrots processed in a peeled baby carrot production process, it is used in sauces, baked goods, bakery mixes, processed meats and other food applications. The e ingredient acts as a binder, thickener, extender and stabilizer and is used at levels not exceeding 5% in the finished product. Given that the product begins with fresh carrot, the extensive processing and extended exposure to air degrades β-carotene extensively to yield OxBC. In a retail sample we have isolated ~0.3 mg carotene copolymer compound/g carrot fiber powder.

Bolthouse Farms, in their National List Petition submission to USDA for use of carrot fiber in organic foods, estimated that the dietary intake of carrot fiber in a single serving is 2.5 g for franks or sausages and 4.2 g for meat patties or canned meat (Bolthouse Farms, 2002, 2007). Using the carotene copolymer content value of 0.3 mg/g fiber as a guide, this translates roughly to ~1 mg OxBC per single item serving. As an indication of the extent of carrot fiber production and its use in the food industry, Bolthouse Farms have stated that carrot fiber is produced in their operation from approximately 100 tons of carrot waste per day (Bolthouse Farms, 2007).

Foods prepared using tomato powder are especially rich in carotenoid copolymer compounds, especially the lycopene copolymer. For example, soups, sauces, stews and casseroles prepared with tomato powder can contain an estimated 52-78 mg of carotenoid copolymer per serving (Supplementary Material S3, Table 2). One serving of a tomato sauce prepared from a published recipe is estimated to provide 62 mg carotenoid copolymer (Supplementary Material S3, Table 3). A recipe for Hungarian goulash using tomato paste and paprika is estimated to provide 20 mg carotenoid copolymer per serving if tomato powder is used. Of note, the lycopene-oxygen copolymer in fully oxidized lycopene is chemically very similar to the β-carotene copolymer (Burton et al., 2016).

#### 3.4.5. Human exposure from livestock fed synthetic OxBC

Assuming a Feed Conversion Ratio (FCR; feed consumed/weight gain) of 3.0 for swine and 1.5 for poultry, respectively, the total amount of OxBC available for uptake from feed can be calculated. As an example, uptake values calculated over the full grow periods for feed containing 4 ppm (i.e., mg/kg) OxBC for swine and 2 ppm OxBC for poultry, which are inclusion levels typically adopted for these species, are 12 mg and 3 mg per kg body weight, respectively. The calculations are as follows: swine, 4 ppm and an FCR of 3.0, 3 kg feed provides 3 x 4 = 12 mg OxBC per kg body weight; broilers, 2 ppm and an FCR of 1.5, 1.5 kg feed provides 1.5 x 2 = 3 mg OxBC per kg body weight.

For a generous serving of 500 g of pork or chicken and assuming muscle is 50% of live body weight, the maximum possible amount of OxBC available to a consumer per serving of meat from full uptake by the animals is 6 mg and 1.5 mg from swine and broiler, respectively. However, incomplete absorption from the gut of the animals, which, for example, for β-carotene is a maximum of 65% in humans (Haskell, 2012), and ongoing metabolism could conservatively reduce the available OxBC by half, i.e., to 3 mg and 1.5 mg per serving for pork and poultry, respectively.

Exposure to synthetic OxBC in milk can also be roughly estimated using the fact that in Ontario a dairy cow produces an average of 30 liters from twice-daily milking (Dairy Farmers of Ontario, 2013). Assuming 300 mg supplementation of OxBC per cow per day, as used in trials (McDougall, 2020), and assuming the daily retention of OxBC is 25% in the milk, the OxBC content in one liter of milk would be 0.25 x 300 mg/30 liter = 2.5 mg/liter.

In foods containing tomato powder, there is a substantially larger level of carotenoid copolymer content, 6-78 mg, arising from the large amount of lycopene originally present in tomatoes (Supplementary Material S3, Tables 1-3).

#### 3.4.6. Exposure to natural OxBC in livestock

OxBC is naturally present in poultry feed, at least in a typical wheat-based feed (Supplementary Material S3, Table 4). The mean value from six feed samples was 1.0 ± 0.4 ppm. It is probable that the naturally occurring OxBC originates from β-carotene originally present in the wheat used in the feeds. The low parts-per-million levels of synthetic OxBC livestock supplementation of 2-4 ppm for poultry feed used by Kang et al. (Kang et al., 2018) and others is comparable to the natural background levels present in wheat-based feeds. Also, the levels of 2-4 ppm of synthetic OxBC used for poultry and 4-8 ppm for swine (Chen et al., 2020; Kang et al., 2018; Kinh et al., 2020) are well within the estimated historical ranges of approximately 2-5 ppm natural OxBC in poultry feed and 3-9 ppm natural OxBC in swine feed, if alfalfa hay or meal were present at 3-7.5% and 5-15%, respectively (Supplementary Material S3, Livestock exposure to natural sources of OxBC).

Alfalfa has a long history as an important forage crop for animals (Supplementary Material S3, Livestock exposure to natural sources of OxBC). Sun-dried alfalfa contains quantities of OxBC and carotenoid copolymers that are a significant fraction of the original carotenoid levels (Burton et al., 2016). When dried into hay, alfalfa is a rich source of OxBC and other carotenoid copolymers, including those of lutein and neoxanthin (Bickoff et al., 1954). It follows that other carotenoid-rich forage crop hays also will contain OxBC and carotenoid copolymer compounds.

A 1940 USDA bulletin entitled “The Uses of Alfalfa” contains information on feeding dried alfalfa to beef cattle, dairy cows, poultry and hogs (Westover and Hosterman, 1940). It was estimated that a dairy cow consumes 9-14 kg of alfalfa hay daily. The value of 61 μg/g (ppm) OxBC in alfalfa from Table 1 in Supplementary Material S3 translates to an intake of 550-850 mg of OxBC daily. This can be compared to the level of 300 mg OxBC/head/d fed to dairy cattle in a trial that showed reductions in sub-clinical mastitis (McDougall, 2020).

Exposure to the more abundant total carotenoid copolymer in alfalfa can be estimated using the value of 978 μg/g dried alfalfa obtained for carotenoid copolymer compounds isolated from alfalfa (Supplementary Material S3, Table 1). A much higher value is obtained because of the original presence of lutein and other carotenoids, in addition to β-carotene. For example, 9 kg alfalfa hay x 978 μg/g alfalfa would provide 8.8 g oxidized carotenoid compounds daily.

## 4. Discussion

### 4.1. Genotoxicity assays

The weak to moderate activity seen in the bacterial reverse mutation assay in the *Salmonella typhimurium* TA 100 strain occurred at the higher concentration range, beginning at 500 μg/plate. This corresponds to an approximate exposure level of 200 ppm OxBC. For comparison the level of exposure in OxBC livestock feeding trials ranges from 2-8 ppm. As some of the small apocarotenoid compounds present in OxBC are potentially chemically reactive electrophiles and carbonyls (Burton et al., 2014; Mogg and Burton, 2020), it is not unexpected that some degree of mutagenicity is seen at higher concentrations. The higher threshold for mutagenicity seen with metabolic activation (i.e., > 500 μg/plate) supports the possibility that metabolism would at least partially mitigate the reactivity of OxBC’s reactive components.

Although OxBC was clastogenic *in vitro* at toxic concentrations in the chromosomal aberration assay, this is not necessarily considered a safety concern, as per ICH S2(R1) (European Medicines Agency, 2013), if an *in vivo* genotoxicity study can be used to establish the genotoxic potential of the compound. Indeed, the acute toxicity test showed a very high tolerance of OxBC in mice (1800 mg/kg) together with a clear negative outcome of the *in vivo* mouse micronucleus assay using a recognized test guideline (OECD; micronucleus assay). Previous dietary supplementation studies with OxBC in cattle and lactating sows provide good evidence of uptake from the gut in the form of systemic biological effects in cattle (Duquette et al., 2014) and appearance of OxBC in sow’s milk (Chen et al., 2020). Evidence of absorption from the gut combined with the findings of the micronucleus assay support that the normally encountered levels of natural dietary OxBC as well as the recommended supplementary levels of synthetic OxBC are well within the safe range for both humans and livestock (Schaub et al., 2017).

Safety in livestock applications is corroborated by the absence of any adverse events observed in the numerous livestock trials conducted during 15 years of low-dose ppm levels of OxBC in feeding trials with thousands of food animals, primarily poultry, dairy cows and swine - including gestating and lactating sows. A history of commercial use of synthetic OxBC as a feed additive in several countries over the past five years provides additional evidence of safety. Synthetic OxBC has been added to an estimated total of 1 million tonnes of feed with no reports of adverse events. Also, no serious adverse events have been reported in monitoring more than one million commercial OxBC administrations to domestic dogs and cats for a decade.

The *in vivo* mouse results strongly suggest that any potentially reactive compounds present in OxBC, such as peroxides, as have been measured in plant food items (Schaub et al., 2017), and reactive carbonyl compounds (Mogg and Burton, 2020), are safely metabolized.

### 4.2. Dietary exposure

OxBC and β-carotene-oxygen copolymers are a normal part of human diets and animal feeds. The close chemical identity of the synthetic OxBC copolymer and the copolymer isolated from carrot powder has allowed assessment of levels of dietary sources of natural OxBC to provide an indirect indication of OxBC safety. Dried forms of carrot and tomato are rich sources of oxidized carotenoids and of the copolymer compounds of β-carotene and lycopene, respectively. The very long history of dietary exposure to these oxidation products provides a measure of support for their safety at their normal levels of dietary intake.

Nowadays, the exposure to these compounds is extensive and widespread in the food industry, especially through use of carrot, tomato and sweet potato powders. The estimated range of exposure to natural OxBC (1-22 mg per serving) is comparable to the level of safe intake of β-carotene (<15 mg/d) that results from the regular consumption of the foods in which β-carotene occurs naturally (5-10 mg/d), in addition to food additives and food supplements (European Food Safety Authority, 2012a). Consumption of common tomato-based products results in exposure to even higher levels of natural oxidized carotenoids (Burton et al., 2016).

Livestock have long been exposed to oxidized carotenoids, especially in the form of forage hays and, in particular, alfalfa. OxBC is naturally present at low levels in poultry feeds, e.g., ~1 ppm in wheat-based feeds. Synthetic OxBC has been used in poultry (2-4 ppm) and swine (4-8 ppm) feeds at levels that are comparable to natural exposures for poultry (2-5 ppm) and swine (3-9 ppm) estimated from historical use of dried alfalfa. Similarly, supplementation in dairy cows (e.g., 300 mg/head/d) is comparable to estimated levels from alfalfa consumption (~550-850 mg/head/d).

For human exposure to synthetic OxBC, the estimated values of 1.5-3 mg for consumption of a single serving of poultry or pork and 2.5 mg from drinking one liter of milk can be compared with the estimated exposure of 1-22 mg of natural OxBC available in a single serving of foods made with carrot powder (Supplementary Material S3, Tables 2 and 3) and are comparable to the estimated ~1 mg level of exposure from a single serving of processed meat containing carrot fiber binder.

The apocarotenoid products in OxBC contain thirteen GRAS human flavor agents and lack the larger, potentially genotoxic, retinoid-like apocarotenoid compounds. Safe use of the many apocarotenoids in synthetic OxBC is supported by the very low ng/g individual levels of these compounds within the low μg/g (ppm) level of OxBC livestock usage, which compares favorably to the ng/g levels of apocarotenoids present in β-carotene-containing plant sources (Schaub et al., 2017).

β-Carotene oxidation is now understood to proceed by competition between polymerization with oxygen, predominantly, to form the β-carotene-oxygen copolymer compound, and depolymerization to form apocarotenoids. The copolymer has been shown to be moderately stable and is the actual source of a minimum of 45 identified apocarotenoids, including GA, and more than 90 other unidentified small molecule compounds.

By studying the biosynthesis and degradation of β-carotene in non-green plant tissue it has been determined that β-carotene degradation occurs mainly by non-enzymatic oxidation to maintain a dynamic balance with biosynthesis (Schaub et al., 2018). However, as shown in various plant food items, apocarotenoids represent only a very small proportion of the degradation products, with the β-carotene-oxygen copolymer being the main product (Schaub et al., 2017) and corroborating our earlier finding regarding the predominance of the natural occurrence of the copolymer. The relative stability of the copolymer suggests that the compound serves not only as an apocarotenoid repository but, importantly, maintains the potentially reactive free apocarotenoids at low concentrations during β-carotene autoxidation (Mogg and Burton, 2020) where, in plants at least, some can serve protective cellular signalling functions (Havaux, 2020).

The genotoxicity results, together with the record of the long-term and extensive dietary exposure to naturally occurring OxBC and the absence of any reports of associated toxicity, support the safety of the natural form and, in particular, the use of closely similar synthetic OxBC in animals at levels comparable to levels typical of natural OxBC. The normally encountered levels of natural OxBC in the diet as well as the recommended inclusion level of the synthetic OxBC-based commercial supplement are several-fold lower than the dosage deemed toxic in the acute toxicity assay, indicating a wide margin of safety for OxBC.

## Supporting information

Genotoxicity methods

Carrot powder oxidation

Dietary exposure to oxidized carotenoids

## Abbreviations

OxBC: fully oxidized β-carotene

## Supplementary material

Supplementary Material S1. Genotoxicity methods

Supplementary Material S2. Carrot powder oxidation

Supplementary Material S3. Dietary exposure to oxidized carotenoids

