## Supplementary material for "β-Carotene oxidation products - function and safety": Genotoxicity methods

### Supplementary Material S1
### Supplementary Materials and Methods

**Bacterial reverse mutation assay (Ames test)**

OxBC was tested for mutagenic activity in *Salmonella typhimurium* strains TA 1535, TA 1537, TA 98 and TA 100 and in *Escherichia coli* WP2*uvr*A. Dimethyl sulfoxide (DMSO) was used as the solvent and vehicle for OxBC with test formulations being prepared immediately prior to dosing (within 1 h). Mutagenic activity was assessed with the bacterial reverse mutation assay (U.S. Food and Drug Administration, 2007). An S9 mixture (S9), a cytosolic homogenate prepared from the livers of Aroclor 1254-treated rats, along with cofactors necessary for enzymatic activity, provided an exogenous metabolic activation system (McGregor et al., 1988).

The four *S. typhimurium strains* were obtained from Professor B. N. Ames at the University of California, USA and maintained in liquid nitrogen storage (Ames et al., 1975). The *E. coli* strain used was WP2*uvr*A, which was obtained from the National Collection of Industrial Bacteria, Aberdeen, Scotland and maintained in liquid nitrogen storage. Samples of each bacterial strain were cultured for 16 h at ~37°C in nutrient broth (25 g Oxoid Nutrient Broth No. 2 per liter). While some of the cultures were kept for up to 7 days at ~4°C to allow relevant checks to be performed, fresh cultures were used in the study tests. Two experiments were conducted in both the absence and the presence of S9. OxBC was dosed at concentrations spaced at half-log intervals in the first mutation assay and at concentrations spaced at halving intervals in the second mutation assay. Triplicate plates were poured for each exposure level (n = 6) and bacterial strain (n = 5) in the absence or presence of S9. A toxicity test was performed to establish the concentration range to be used in the first mutation test, covering 17, 50, 167, 500, 1667 and 5000 µg per plate. The concentrations tested in the first mutation experiment in the absence of S9 were as follows for all 5 bacterial strains: 5, 17, 50, 167, 500 and 1667 µg per plate. In the presence of S9, the dose levels selected for all 5 bacterial strains were: 17, 50, 167, 500, 1667 and 5000 µg per plate. In the second mutation experiment the concentration levels depended on the first mutation experiment results. Six concentrations were tested, spaced at halving intervals. These were, in the absence of S9, for all 5 bacterial strains: 78, 156, 313, 625, 1250 and 2500 µg per plate, and in the presence of S9, for all 5 bacterial strains: 156, 313, 625, 1250, 2500 and 5000 µg per plate. DMSO was plated in triplicate for each strain used, in the presence and absence of S9.

The following positive control concentrations were plated in triplicate with S9: 2-aminoanthracene (2AAN): 2 µg per plate with *S. typhimurium* TA 1535 and TA 1537, 0.5 µg per plate with *S. typhimurium* TA 98 and TA 100 and 20 µg per plate with *E. coli* WP2*uvr*A; Without S9: sodium azide ($NaN_3$): 1 µg per plate with *S. typhimurium* TA 1535 and TA 100; N-ethyl-N-nitro-N-nitrosoguanidine (ENNG): 2 µg per plate with *E. coli* WP2*uvr*A; 2-nitrofluorene (2-NF): 1 µg per plate with *S. typhimurium* TA 98; 9-aminoacridine (9-AA): 80 µg per plate with *S. typhimurium* TA 1537. All OxBC concentrations were plated in triplicate.

*S9 Mixture:* S9 enzymes, prepared in-house from livers of Aroclor-1254-treated adult, male Fischer rats were stored in sterile plastic tubes immersed in liquid nitrogen (*ca* -196°C) and used within 6 months of preparation. The enzymatic activity of each batch was characterised by testing selected pre-mutagens in an Ames test with *S. typhimurium* TA 1538. Although strain TA 1538 was not used in this study, considerable historical data exists on enzymatic activity with this *S. typhimurium* strain, and these data were used for comparison with current batches of S9 enzymes.

*Toxicity Test.* To establish suitable exposure levels for the first mutation test, an initial dose-finding test was conducted in the presence and absence of S9. A single strain of *S. typhimurium* TA 100 was used, and one plate per exposure level of OxBC was prepared. Plates were incubated for 2 days, the numbers of revertant colonies were then noted and the plates carefully examined, microscopically, for thinning of the background lawn of microcolonies. Condition of background lawn was assessed as normal, slightly thin lawn (ST), thin lawn (TL), very thin lawn (VT) or lawn absent (A). Any precipitation of OxBC on the plates was noted.

*Mutation Tests.* Diluted agar (0.6% Difco Bacto-agar, 0.6% NaCl) was autoclaved and, just before use, supplemented as follows: For *S. typhimurium,* a sterile 1.0 mM L-histidine·HCl, 1.0 mM biotin solution was added at 50 mL per litre of soft agar. For *E. coli*, a sterile 1.35 mM L-tryptophan solution was added at 10 mL per litre of soft agar. The soft agars were thoroughly mixed and kept in a water bath at 45°C.

*Treatment Method (Direct Plate).* Two mL of soft agar were dispensed into a small, plastic, sterile tube. Either S9 or 0.05 M phosphate buffer, pH 7.4 (0.5 mL) was added, followed by 0.1 mL bacteria and, finally, the OxBC solution (0.1 mL). The tube contents, which were continuously cooled, were mixed and then poured onto minimal medium plates that had been prepared in-house. The minimal medium plates contained 20 mL of 1.5% BBL purified agar in Vogel-Bonner Medium E (Vogel and Bonner, 1956) with 2% glucose. After the soft agar had set, plates were inverted and incubated at 37°C for 3 days and then examined. The number of mutant colonies on each plate were determined using a Sorcerer Colony Counter and captured electronically in a validated software system Ames Study Manager. The plates were also examined microscopically for precipitates and for microcolony growth.

*Quality Control of Bacterial Strains.* All bacterial strains were plated, one plate per strain, onto complete medium and tested for ampicillin resistance and crystal violet sensitivity. All bacterial strains were also checked for UV-radiation sensitivity and for essential amino acid requirements.

*Calculations and Data Acceptance.* The mean number of mutant colonies, plus standard deviation, was calculated for each set of three plates. In addition, the fold-increase over the vehicle control was calculated for all test item and positive control treatments. Historical vehicle and positive control values were used to assess the acceptability of the results. A test was acceptable if the following occurred: Each bacterial strain demonstrated typical responses to crystal violet, ampicillin and ultraviolet radiation. At least 2 of the 3 vehicle control plates were within the following ranges for mean number of revertant colonies: *S. typhimurium* TA 1535: 4-30; TA 100: 60-200; TA 1537: 1-30; TA 98: 10-60 and *E. coli* WP2*uvrA*: 1-60. There were at least 3-fold increases over the mean vehicle control values in at least 2 of the 3 positive control plates for each strain and activation state (in the case of TA 98 and TA 100, at least 2-fold was required). No toxicity or contamination was observed in at least four concentration levels.

*Interpretation of Mutagenicity.* For *S. typhimurium* strains TA 1535 and TA 1537 and for *E. coli* WP2*uvr*A, at least a 3-fold increase over the mean concurrent vehicle control value was required before mutagenic activity was suspected. For *S. typhimurium* strains TA 98 and TA 100, a 2-fold increase over the control value was considered indicative of a mutagenic effect. For *S. typhimurium* strains TA 1535 and TA 1537 and for *E. coli* WP2*uvr*A, a minimum count of 20 was required before a response was identified. A concentration-related response was also required for identification of a mutagenic effect. At high concentrations, this relationship may be reversed, due to, for example, toxicity of the test item to the bacteria, specific toxicity of the test item to the mutants, or inhibition of S9 enzymes (where a mutagen requires metabolic activation by S9). No statistical analysis was performed.

**Chromosomal aberration assay**

The assay tested the ability of OxBC to induce chromosomal aberrations in cultured Chinese hamster ovary cells. The study complied with OECD and ICH Guidelines, and with the European Commission Annex V, Test Method B10. DMSO was used as the solvent and vehicle for OxBC throughout the study. The CHO 10 B4 Chinese hamster ovary cells, routinely tested for mycoplasma, were grown as monolayers with a generation time of ~12 h. The modal chromosome number determined for these cells is 21. Cells were incubated at 37°C in a basic medium (Ham's F-10) containing HEPES buffer, supplemented with the antibiotic minocycline. For cell growth and treatment in the absence of S9, fetal bovine serum (10% v/v) was added. The medium used for treatment in the presence of S9 and for washing cultures before or after treatment was serum free. Cells were trypsinized from stock flasks at passage numbers 16 (Toxicity Test) and 17 (Test 1) and resuspended in fresh culture medium at a density of 0.1 x $10^6$ cells/mL. The cells, in 5 mL volumes, were dispensed into 25 $cm^2$ tissue culture flasks.

Test cultures were established from the stock flask ~20 h before testing. For the toxicity test, 9 dose levels were tested. The highest dose was 5000 μg/mL, the maximum allowable concentration, and subsequent dose levels were halving dilutions. In Test 1, the dose levels in the presence of S9 were: 20, 40, 50, 60, 70, 80, 90 and 100 μg/mL; and in the absence of S9 were: 10, 20, 30, 40, 50, 70, 80 and 90 μg/mL. All experiments included vehicle control cultures which were subjected to the same experimental manipulations as treated cultures, both in the presence and absence of S9. The following positive control materials were used: *with S9* - cyclophosphamide 20-50 μg/mL; *without S9* - methyl methanesulfonate 10-40 μg/mL. The enzymic activity of each batch of S9 was characterised by testing selected pre-mutagens in an Ames test with *S. typhimurium* TA 1538. S9 batches used demonstrated, within each test, a satisfactory clastogenic response in cells treated with cyclophosphamide.

Precipitation was noted in cultures treated with 156-5000 μg/mL OxBC in the presence of S9 and in cultures treated with 313-5000 μg/mL in the absence of S9 (6 h treatment) and 625-5000 μg/mL (22 h treatment) during the toxicity test. In the main aberration assay, cultures were treated up to a concentration of 100 μg/mL in the presence of S9 and up to a concentration of 90 μg/mL in the absence of S9. Precipitation was noted in the cultures treated with 80-100 μg/mL in the presence of S9, while no precipitation was noted in the absence of S9. OxBC did not change the colour of the culture medium, therefore no pH measurements were made. Cultures were established after 20 hours of pre-exposure in the presence or absence of S9. For both the toxicity test and Test 1, the schedule was as follows: treatment, 0-6 h; recovery, 6-22 h; colcemid (if required), 22-24 h; harvest, 24 h.

Treatments with OxBC or the vehicle control were performed on single cultures in the toxicity test and in duplicate cell cultures during Test 1. Several concentrations of the positive controls were tested using single cultures in Test 1. Cultures treated in the presence of S9 were washed before treatment with serum-free medium, and exposure medium was prepared, immediately before dosing, in sterile containers. The dosing solution (50 µL) was administered to cultured cells with or without 0.5 mL S9. After treatment, cells were washed twice with serum-free medium, then growth medium (and colcemid, if required) was added at 4.5 mL with S9 or 5.0 mL in its absence for the recovery period to a final volume of 5 mL. Living cultures were examined for evidence of changes to cell morphology, once at the end of the treatment period and again before harvesting. In Test 1, colcemid was added to all cultures to a final concentration of 0.1 µg/mL. Cells were cultured in medium containing colcemid for 2 h and then accumulated in metaphase. In Test 1, mitotic cells were harvested by gently tapping flasks to release them from the monolayer. Cells were sedimented by centrifugation (~190 *g*) and treated with hypotonic solution (1% trisodium citrate) for 15 min at room temperature. The cells were then fixed (after sedimentation) using 4 mL freshly prepared fixative (methanol:glacial acetic acid, 3:1). Two further changes (after sedimentation) of fixative were made. In the toxicity test and in Test 1, monolayer cells were trypsinized, counted and discarded. This provided a quantitative measure of toxicity. For Test 1, three slides per culture were prepared. Slides were prepared by dropping the cell suspension onto clean, grease-free slides. The slides were stained with 5% Giemsa and then made permanent by mounting coverslips with DPX mountant. The slides were then examined for evidence of metaphase cells and signs of cellular necrosis. Based on toxicity (i.e., cell counts and slide/culture observations) and osmolality, 3-4 concentration levels were selected for assessment of chromosomal aberrations. From 2-3 slides per culture and up to 50 metaphase cells per slide, a total of 100 metaphase cells per culture were examined, where possible. A reduced number of metaphases were scored if a high proportion (≥40%) of metaphase cells were found to be damaged. A microscope was used for this assessment at a magnification of *x 1000* or *x 1250*. The number of chromosomes in each metaphase cell and all abnormalities, using the nomenclature of Gebhart (Gebhart, 1970), were recorded.

*Quantitative Measure of Toxicity.* The number of cells recovered per culture was calculated from the cell counts and this was then compared with the number of cells (the mean of two cultures) recovered from the vehicle control cultures. Four parameters were calculated and judged as negative, suspicious or positive. These parameters were: i) lesions per cell, ii) percentage of aberrant cells, including cells with gaps only, iii) percentage of aberrant cells, excluding cells with gaps only, and iv) percentage of aneuploid cells. The third parameter is considered to be the most important of the four in judging the true clastogenicity of a test item. The results obtained were compared with historical control data.

A dose level was considered to be toxic if the cell count was reduced to <50% of the mean vehicle control culture values or if consistent evidence of changes to cell morphology was observed. The results for OxBC and positive control treated cultures were evaluated by comparison with the concurrent vehicle control cultures and with historical negative control data. A negative response was recorded if responses from the test item treated cultures were within 95% confidence limits for the historical negative control data. The response at a single dose was classified as significant if the percent of aberrant cells was consistently greater than the 99% confidence limits for the historical negative control data or greater than double the frequency of an elevated vehicle or untreated control culture, if appropriate. A test was positive if the response in at least one acceptable dose level was significant by the criterion described above. A test item was positive if Test 1 was positive, as described above, or if one of the tests was positive and the other test indicated activity. An indication of activity could be suspicious levels of aberrant cells (between 95% and 99% confidence limits). Experiments that met in part the criteria for a positive response, or marginally met all the criteria, were classed as inconclusive. No statistical analysis was performed.

**Mouse micronucleus assay**

OxBC was evaluated for *in vivo* clastogenic activity and/or disruption of the mitotic apparatus by detecting micronuclei in polychromatic erythrocytes in CD-1 mouse bone marrow. Experimental procedures complied with OECD, ICH and EC Guidelines, US EPA Pesticide Assessment Guidelines, and the Japanese Guidelines on Genotoxicity Testing and recommendations published by the US EPA Gene-Tox Program and the Japanese Collaborative Study Group for Micronucleus Testing (Environmental Mutagen Society, 1990; Mavournin et al., 1990).

The vehicle control material used for OxBC was a solution of N-methyl-2-pyrrolidone, polyethylene glycol 400 and propylene glycol prepared at Charles River in the ratio of 1:2:2 respectively. The OxBC dose volume used for both the control and test item treated animals was a constant 10 mL/kg body weight. Cyclophosphamide was prepared fresh as a 5 mg/mL solution in water, and it was administered to the positive control animals in dose volumes of 10 mL/kg to provide the required target dose of 50 mg/kg. Male and female CD-1 mice were supplied by Charles River UK, (Kent, England). All mice were 6-7 weeks of age at the time of dosing. The toxicity test was conducted in a stepwise manner using 3 phases to determine the maximum tolerated dose of OxBC. The starting dose level was 2000 mg/kg, which seemed acceptable, however, a further toxicity test at 2000 mg/kg deemed this dose level as unsuitable. A subsequent test at 1600 mg/kg/day showed this dose level was not sufficient to be classified as the Maximum Tolerated Dose. The mice were observed for clinical signs or mortality at frequent intervals (1 min, 0.5 h, 1 h, 2 h and 4 h) post-dosing, and then twice daily until the end of the observation period. No difference in toxicity was observed between the male and female groups, therefore only male mice were used for the micronucleus test, and they were dosed at 1800 mg/kg/day.

In the micronucleus test, groups of mice were dosed orally at 0 h and 24 h with test or control materials, and marrow samples were taken 24 h later as follows (dose group, daily test dose, number of mice): Vehicle Control, 10 mL vehicle/kg, 5 M; Low Dose, 450 mg OxBC/kg, 5 M; Mid Dose, 900 mg OxBC/kg, 5 M; High Dose, 1800 mg OxBC/kg, 9 M (one animal died due to technical problems prior to dosing); Positive Control, 50 mg cyclophosphamide/kg, 5 M; Untreated Control, 3 M. Animal health status checks were done at frequent intervals after dosing and prior to the scheduled kill. Mice were killed by cervical dislocation. One femur from each mouse was quickly dissected out and freed of adherent tissue. A small hole was made in the neck of one femur, and the marrow was flushed using a 1 mL syringe fitted with a 25-gauge needle, into a centrifuge tube containing ~3 mL of a 1:1 mixture of fetal calf serum and 0.8% trisodium citrate in Sorensen's buffer, pH 6.8 (Sorensen's buffer, pH 6.8, 2.84 g $Na_2HPO_4$/L plus 2.72 g $KH_2PO_4$/L distilled water). Routine tissue culture antibiotics were included to prevent microbial growth. This mildly hypotonic treatment served to make the micronuclei clearly visible and to distinguish them from surrounding artefacts. Following completion of the sampling procedure, the contents of the tubes were briefly agitated on a vortex mixer to allow for cell separation, and the tubes were centrifuged to pellet the cells. All but a few drops of supernatant fluid were discarded. The cells were then resuspended on a vortex mixer in the residual amount of supernatant liquid. A few drops of the suspension were placed at one end of the slide, and a smear was made by drawing the top of a glass Pasteur pipette horizontally along the slide. Two slides were prepared from each tube per animal. The smear was left to air dry, fixed in methanol for ~5 min, immersed for ~15 min in 15% Giemsa stain, and then prepared in water to give optimum erythrocyte discrimination. The stained smears were finally rinsed in distilled water for 1 min and left to air dry overnight. Permanent slide preparations were made by sealing coverslips onto the glass slides using DPX mounting medium.

The better of the two prepared slides was selected for examination, and the coded slides were assessed blind by the same operator. At least 2000 polychromatic erythrocytes (PCE) per animal were scored for micronuclei, and the frequency of micro-nucleated cells (MN-PCE) was determined. As a control against the inclusion of artefacts or the action of a mutagen on the $G_2$ and/or mitotic phase of the cell cycle, the numbers of micro-nucleated normo-chromatic erythrocytes (MN- NCE) in mature red blood corpuscles were also recorded (Hamoud et al., 1989; Maier and Schmid, 1976; Schmid, 1976). In addition, scored micronuclei were assigned on the basis of size into small or large categories, historically defined as micronuclei occupying less or more than 25% of the visible cellular area. This classification provided a non-specific measure of compound-induced spindle dysfunction, as large micronuclei appear to derive from lagging chromosomes caused by damage to the mitotic apparatus during bone marrow erythropoiesis (Vanderkerken et al., 1989; Yamamoto and Kikuchi, 1980). The PCE/NCE ratio, a measure of any induced systemic toxicity, was determined by counting a minimum total of 1000 erythrocytes (PCE plus NCE) per marrow preparation. The scoring was done under a nominal magnification of *x 1250*.

The following three acceptance criteria were satisfied: i) prepared slides had uniform staining properties and a sufficient number of PCE cells present to allow accurate micronucleus determination, ii) the assay was considered acceptable as the MN-PCE frequencies for the vehicle control dosed mice were within the expected historical range, with the ranges being defined in accordance with Charles River experience with the bone marrow micronucleus test using CD-1 mice, and iii) there was an adequate positive control response for at least 2 animals and the dose group as a whole.

The average micronucleus incidence in vehicle control-dosed and untreated CD-1 mice, had been determined at the laboratory as 0.05±0.06%, a range of 0.00-0.23% per group of 5 animals, in agreement with published data for micronucleus tests with CD-1 mice (Salamone and Mavournin, 1994; Tamura et al., 1990). These historical data were used in the response evaluation for this test. The test was judged negative if no biologically relevant increases in the numbers of MN-PCE were observed, relative to the concurrent and established historical control frequencies for MN-PCE induction. No statistical analysis was performed if the levels of MN-PCE induction fell within the determined historical control frequencies. A similar biological approach to the data, which avoids the need for statistical evaluations was described by Ashby and Tinwell (Ashby and Tinwell, 1995). Variations in the MN-NCE frequencies and PCE/NCE ratios were also not analysed statistically, unless clearly different from concurrent control values. The test was judged positive if an increase in the number of MN-PCE was obtained for one or more of the OxBC-treated dose groups. That is, an increase >10% over the expected historical control ranges for a group of animals. The increase observed needed to be biologically relevant and statistically significant relative to concurrent and historical control frequencies for MN-PCE and/or MN-NCE induction. The test was considered inconclusive if the levels of MN-PCE within any one dose group were increased above the established historical control frequencies for MN-

PCE induction, but not high enough to meet the criteria for a positive response (i.e., an increase up to 10% over the maximum negative control frequency for a group of animals).
