## Supplementary material for "β-Carotene oxidation products - function and safety": Carrot powder oxidation

### Supplementary Material S2
### β-Carotene Oxidation in Carrot Powder

#### Introduction

This study illustrates the importance of surface area, dehydration and exposure to air in enhancing β-carotene oxidation through the example of puréeing of carrot followed by dehydration and powdering. Oxidation in the carrot was followed by monitoring the decrease in β-carotene concentration and the simultaneous increase in geronic acid concentration. Geronic acid is an indirect marker of OxBC and associated carotenoid-oxygen copolymer formation (Burton et al., 2016; Burton et al., 2014).

#### Results and Discussion

Fresh, shredded carrots were found to contain 4.9 ng/g geronic acid (Table 1). Dehydration of the carrot purée resulted in a decrease in mass of 89.5%. As seen in Table 1, the amount of geronic acid in the processed carrot after 1 day (Day 0) had increased approximately 20-fold after puréeing and dehydration. The increased level of geronic acid is a result of both a 10-fold decrease in mass during dehydration and oxidation. Further processing by powdering the dried purée and spreading it thinly on a tray to expose it to air and light caused an approximate 9-fold increase in geronic acid after 5 days, with an accompanying decrease in β-carotene. After 21 days, geronic acid had increased substantially to 4155 ng/g, and β-carotene had decreased to less than half its amount at Day 0. The calculated levels of OxBC are shown in Table 1.

Fig. 1 shows graphically the inverse relationship of the rise of geronic acid to the decrease of β-carotene. Dividing the change in OxBC by the change in β-carotene at each time point after time 0 gives a ratio that is roughly constant (Table 1), indicating that production of geronic acid and. indirectly, OxBC is closely linked to the oxidative loss of β-carotene.

**Table 1**. Effect of dehydration upon β-carotene and geronic acid levels in carrots.

| Carrot Physical State | Time (days) | β-carotene[a] (μg/g) | Geronic Acid (ng/g) | OxBC (Calc)[b] (μg/g) | Δ β-carotene/ Δ OxBC |
|---|---|---|---|---|---|
| Fresh | -1 | n.d.[c] | 4.9 ± 1.9 | 0.34 | - |
| Purée, dried | 0 | 1366 (99.5%) | 97 ± 18 | 6.8 | - |
| Powder, dried | 5 | 1141 (95%) | 879 ± 56 | 62 | 4.1 |
| Powder, dried | 12 | 697 (74%) | 3478 ± 252 | 243 | 2.8 |
| Powder, dried | 21 | 573 (66%) | 4155 ± 22 | 291 | 2.8 |

[a] The β-carotene measurement is approximate. The assay measures the absorbance of whole carrot extract at 454 nm, the maximum absorbance wavelength of β-carotene. However, there are smaller amounts of other compounds present, such as α-carotene and partially oxidized β- and α-carotenes that could contribute modestly to the absorbance at this wavelength. β-Carotene as a percentage of the sum of β-carotene and OxBC is given in parentheses.
[b] OxBC content calculated by multiplying geronic acid by 70 using an estimated 1.4% geronic acid content in OxBC.
[c] n.d. - not determined

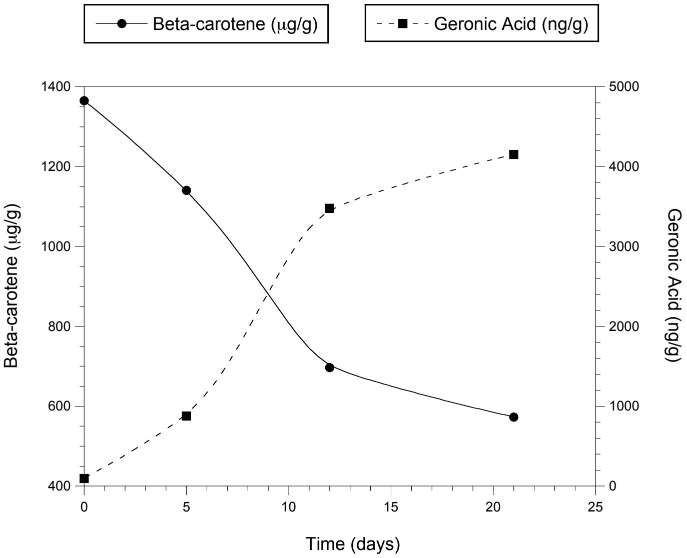


**Figure 1**. Formation of geronic acid and associated loss of β-carotene in dehydrated carrot. Measurements at time 0 used freshly dehydrated carrot purée, and subsequent time points were measured with dried carrot powder, spread thinly on a tray and exposed to air and light.

### Experimental

Fresh, peeled carrots with tops and bottoms cut off were rinsed with water, patted dry with a paper towel and finely shredded with a food processor. Carrot shreds were analyzed for geronic acid to determine the content in fresh carrot. Shreds were also puréed using a food processor, spread approximately ¼ inch thick onto parchment paper, dehydrated in a food dehydrator (Excalibur 3926TB) for 10 hrs at ca. 52-53°C, then allowed to cool to room temp for ca. 8 hrs. The dried purée was analyzed for geronic acid and β-carotene content. The remaining dried purée was powdered using a food processor blade then sifted through a kitchen sieve to remove large particles. The carrot powder was spread thinly onto a tray (46 cm x 36 cm) lined with aluminum foil and placed in an open space. Illumination with a fluorescent light approximately 1.1 meters above the tray was used to approximately simulate exposure to ambient light conditions. Samples of carrot powder were taken at intervals 5-9 days apart for analysis of geronic acid and β-carotene content. Before taking samples of the powder, the tray was gently shaken to mix the powder on the top with that lying beneath it.

***Geronic Acid Assay of Fresh Carrot***

Fresh, peeled carrots with tops and bottoms cut off were rinsed and patted dry with paper towel, then finely shredded using a food processor. 230 - 400 g of carrot shreds were placed in a 1 L beaker, followed by 0.4 mL of geronic acid-$d_6$ solution (0.001604 mg/mL in methanol), BHT (butylated hydroxytoluene; 4-5 mg), and $CHCl_3$ (400-500 mL). The mixture was homogenized for 20 min at 13,500 rpm, then the homogenizer was shut off and the liquid allowed to drain into the beaker, rinsing the homogenizer inside and outside with $CHCl_3$ (total of 4 x 1.5 mL). All material was transferred to a 2 L separatory funnel, the orange $CHCl_3$ layer was separated from the orange pulp, and the pulp was returned to the beaker. $CHCl_3$ (300-400 mL, containing 2-4 mg BHT) was added to the pulp and homogenized for 5 min. The $CHCl_3$ layer was separated as before, the $CHCl_3$ extracts were combined, and solvent was evaporated on the rotavap. $CHCl_3$ was added (15 mL) to dissolve the residue, and the mixture was dried ($MgSO_4$) and filtered through cotton, rinsing with $CHCl_3$ (total 15 mL). Aqueous KOH (25 mL; prepared by dissolving ~ 90 mg KOH in 50 mL water) was added to the $CHCl_3$ solution and stirred vigorously for 5 min. The mixture was transferred to a separatory funnel, most of the $CHCl_3$ was removed, and the remaining liquid was centrifuged for 5 min. The aqueous layer was separated, acidified (~ 3 mL of 1 M HCl), and extracted with $CHCl_3$ (2 x 15 mL). The combined extracts were dried ($Na_2SO_4$), filtered and solvent evaporated. The residue was dissolved in methanol (9 mL), solid $NaHCO_3$ (~ 0.1 g) was added and the mixture stirred. Aqueous $NaHCO_3$ (1 mL, 1 M) was added, followed by $Me_3OBF_4$ (5 small spatulas, ~ 0.3 g). After 15 min stirring, 9 mL $H_2O$ was added, stirred, and the mixture extracted with $CH_2Cl_2$ (2 x 7.5 mL). The combined $CH_2Cl_2$ extracts were dried with $Na_2SO_4$, filtered through cotton and solvent carefully evaporated on the rotary evaporator at room temperature. The residue was dissolved in 0.2 mL acetonitrile and 1 μL injected into the GC-MS (splitless injection, SIM mode monitoring ions 154.1 and 160.1).

***Geronic Acid Assay of Dried Carrot Purée or Powder***

All ethyl acetate used in this procedure contained 0.05 mg/mL BHT. Approximately 3.5 g of dried carrot purée or powder was weighed in a 50 mL test tube (carrot purée was crushed with a spatula to fit it into the bottom of the tube). To the tube was added 15 mL ethyl acetate and geronic acid-$d_6$ (0.64 – 13 μg in methanol). The mixture was homogenized for 10 min at 13,500 rpm, then 10 min at 6500 rpm. The homogenizer was shut off and the liquid allowed to drain into the tube, rinsing the homogenizer inside and outside with ethyl acetate (total of 4 x 1.5 mL). All material was transferred to 2 x 15 mL centrifuge test tubes and centrifuged (~ 5 min). The supernatant was transferred to a 50 mL round bottom flask and solvent was evaporated. At this point the flask was sealed under argon and stored overnight in the freezer until the next morning. Then, 6 mL of 5% aqueous $NH_3$ and 3 mL $H_2O$ were added and stirred vigorously for 20 min. In the meantime, SPE cartridges (Waters Oasis MAX, 500 mg / 6 mL) were prepared by passing through the following solutions in sequence: methanol (6 mL), $H_2O$ (6 mL), 0.5% aqueous $NH_3$ (4.5 mL). The basic solution of analyte was then passed through the cartridge by gravity (pressure with a pipette bulb is acceptable if the flow is very slow). The cartridge was then washed with 0.5% aqueous $NH_3$ (4.5 mL), followed by methanol (9 mL). Carboxylic acids were then eluted from the cartridge by passing through a solution of 2% HCl in methanol (4.5 mL) and collected in a 20 mL scintillation vial.

Solid $NaHCO_3$ was added and stirred until bubbling ceased (~ 30 sec). Then, 1 M $NaHCO_3$ (1 mL) was added, giving a cloudy solution. The solution was stirred gently while $Me_3OBF_4$ was added (5 small spatulas, ~ 0.3 g). The mixture was stirred vigorously for 15 min and maintained slightly basic by ensuring the presence of a small amount of solid $NaHCO_3$ in the vial (visual inspection – adding more if necessary). After 15 min, 6 mL $H_2O$ was added, stirred, and the mixture extracted with $CH_2Cl_2$ (2 x 6 mL). The combined $CH_2Cl_2$ extracts were dried with $Na_2SO_4$, filtered through cotton and the solvent carefully evaporated on the rotary evaporator at room temp. The residue was dissolved in 0.2 mL acetonitrile, filtered, if necessary, through a 0.2 µm Teflon syringe filter, and 1 µL injected into the GC-MS (splitless injection, SIM mode monitoring ions 154.1 and 160.1).

***β-Carotene Assay of Dried Carrot Purée or Powder***

All ethyl acetate used in this procedure contained 0.05 mg/mL BHT. Approximately 1.0 g of dried carrot purée or powder was weighed in a 50 mL test tube. To the tube was added 20 mL ethyl acetate. It was homogenized for 10 min at 13,500 rpm, then the homogenizer was shut off and the liquid allowed to drain into the tube, rinsing the homogenizer inside and outside with ethyl acetate (5 x 1.5 mL). All material was transferred to 2 x 15 mL centrifuge test tubes, centrifuged for ~ 5 min, and the supernatant transferred to a 100 mL volumetric flask. The residue was transferred back to the 50 mL test tube, rinsing the centrifuge tubes as needed with ethyl acetate to ensure complete transfer. A total of 20 mL ethyl acetate was added to the residue and homogenized for 3 min at 6500 rpm. The mixture was centrifuged and separated as before, and the residue extracted once more (3 min, 6500 rpm). The liquid from all 3 extractions were combined into the 100 mL volumetric flask, diluted to volume with ethyl acetate and inverted 30 times to mix. 1 mL of this cloudy orange solution was filtered through a 0.2 µm Teflon syringe filter into a 1 mL volumetric vial. The solution was transferred by pipette to a 10 mL volumetric flask, rinsing carefully with several portions of ethyl acetate to ensure complete transfer. The solution was made up to the 10 mL mark with ethyl acetate, inverted 30 times to mix and the absorbance of the solution was measured at 454 nm. Using the previously determined β-carotene extinction coefficient of 237.67 $mL \cdot mg^{-1} \cdot cm^{-1}$, the amount of β-carotene in solution was calculated.
