## Supplementary material for "β-Carotene oxidation products - function and safety": Dietary exposure to oxidized carotenoids

### Supplementary Material S3
### Exposure to Oxidized β-Carotene (OxBC) and other Oxidized Carotenoids

Abbreviations: GA, geronic acid; OxBC, Oxidized β-carotene

**Background**

Carotenoids are ubiquitous in nature and present in a wide variety of commonly consumed food products. Given their natural tendency to degrade in air, carotenoid oxidation products also are ubiquitous. Whereas the many carotenoid cleavage products (apocarotenoids) are already known, many of them for their flavours and aromas (Winterhalter and Rouseff, 2001), awareness of the existence and immunological activity of carotenoid-oxygen copolymer degradation products happened only recently (Burton et al., 2014; Johnston et al., 2014).

A model oxidation carried out with pure synthetic β-carotene and oxygen in ethyl acetate solvent showed that β-carotene-oxygen copolymers form spontaneously as the predominant degradation product in oxidized β-carotene (OxBC) (Burton et al., 2014). It also was established that the low molecular weight cleavage compound, geronic acid (GA), is a suitable marker for the formation of OxBC and the difficult to detect and measure β-carotene copolymer compounds.

It was anticipated β-carotene would degrade similarly in plant products exposed to air during drying or storage.

We reported the natural occurrence of β-carotene copolymers and other carotenoid copolymers throughout the plant world, especially in the form of fruits and vegetables (Burton et al., 2016). Use of GA confirmed the existence and extent of β-carotene oxidation and provided estimates of the level of OxBC in numerous β-carotene-rich, plant-based foods, as well as certain foods such as milk and eggs derived from animals fed a plant-based diet. The results for a broad array of items are shown in Table 1, updated from the earlier published values (Burton et al., 2016), with OxBC values modestly increased to more accurately reflect the lower level of GA present in OxBC that is formed by air oxidation.

It is apparent in Table 1 that dried plant foods can have an abundance of GA, which inevitably occurs concurrently with substantial associated β-carotene loss. This is illustrated by the two samples of carrot powder, which are distinguished by their colors, orange and light brown, respectively, reflecting their relative β-carotene contents. The light brown carrot powder with no measurable β-carotene has almost double the level of GA and estimated OxBC compared to the orange powder, which still has roughly one quarter of its original β-carotene. Also, the estimated levels of OxBC are in good agreement with the corresponding quantities of isolated copolymer compounds.

The inverse relationship between the disappearance of β-carotene and the appearance of GA has been clearly demonstrated in a model study of the formation of carrot powder from carrot purée (Supplementary Material S2, Table 1, Fig. 1).

The Beyer research group independently has corroborated that β-carotene copolymer compounds are the predominant β-carotene degradation product in their analysis of over 100 plant items (Schaub et al., 2017).

Relatively large amounts of carotenoid copolymer-containing products also were isolated from dried foods in which carotenoids other than β-carotene are abundant (e.g., lycopene, lutein, and capsanthin) (Table 1; this is updated from an earlier version (Burton et al., 2016)). These foods include tomato powder, rosehip powder, paprika, sun-cured alfalfa, and wheatgrass powder. In several dried foods, the level of copolymers is comparable to the original level of the parent carotenoid (e.g., carrot and tomato powders).

Synthetic OxBC, prepared by the full oxidation of pure synthetic β-carotene, has no residual β-carotene nor vitamin A and lacks vitamin A activity. The substance has been shown to both enhance innate immune function (Johnston et al., 2014) and dampen or help resolve excess inflammation (Duquette et al., 2014). The available evidence points to the β-carotene copolymers as the agents responsible for supporting immune function (Johnston et al., 2014).

In livestock, synthetic OxBC added at low μg/g (ppm) levels to feed has shown clear health benefits in various trials with poultry (Kang et al., 2018), swine (Hurnik et al., 2011) and dairy cattle (COFCO, 2018; McDougall, 2020). Enhancing immune function supports better health, which translates into better productivity.

Various health benefits, including improved coat and mobility, consistent with immune function support, have been observed in companion animals, especially canines.

The discovery that the β-carotene copolymer compounds in OxBC have beneficial, non-vitamin A immunological activities led us to surmise their counterparts in foods may impart bioactivities with significant health implications. In humans, carotenoid copolymers could contribute to the beneficial health effects associated with fruit and vegetable consumption (Peto et al., 1981). *In situ* loss of dietary β-carotene resulting from oxidative processes during digestion of fruit or vegetables could generate OxBC and also, at least partially, account for the variable vitamin A activity of β-carotene in foods (Tang and Russell, 2009). Although oxidative destruction of β-carotene will cause loss of vitamin A activity, copolymer formation nevertheless could provide beneficial immunological support.

The knowledge that both OxBC and carotenoid copolymer levels exist in a variety of dried plant products has made it possible to estimate current and historical levels of dietary exposure to these compounds in foods and feeds. This exercise also is of relevance because in 1958 the U.S. Food, Drug and Cosmetics Act was amended by the addition of Section 409, which required the FDA to approve new food additives before they could be used in foods (FDA). Under sections 201(s) and 409 of the Act, and FDA's implementing regulations in 21 CFR 170.3 and 21 CFR 170.30, the use of a food substance may be GRAS either through scientific procedures or, for a substance used in food before 1958, through experience based on common use in food.

**Ubiquity of oxidized β-carotene and the β-carotene-oxygen copolymer**

The work of the Avivagen and Schaub-Beyer research groups independently established that β-carotene copolymers are commonly found in a variety of foods and feeds, including cereals, green and non-green vegetables and fruits, that are consumed regularly by humans and animals.

Schaub and coworkers (Schaub et al., 2017) identified and quantified β-carotene apocarotenoid cleavage products from 102 food items native to the Philippines as well as unprocessed field-grown orange corn cobs and yellow cassava storage roots from Zambia and field-grown orange fleshed sweet potato tubers from Uganda. Their data revealed the ubiquitous occurrence of all possible direct cleavage products and of highly oxidized β-carotene copolymers in plant sources regardless of whether they were biofortified with β-carotene or not.

The motivation behind this work with a Philippine "grocery basket" of products was two-fold: 1) the global problem of vitamin A deficiency, particularly in undeveloped countries, where the deficiency can lead to vision problems, immune-deficiency, and impaired growth (Akhtar et al., 2013; Solon et al., 1978) and 2) questions in conjunction with negative outcomes for smokers and asbestos-exposed populations in the CARET (Goodman et al., 2004) and the ATBC (Virtamo et al., 2014) clinical trials, in which high doses of β-carotene were supplemented and subsequently certain β-carotene cleavage products were suggested as the putative harmful agents (Eroglu et al., 2012).

For these reasons, a wide-variety of fruits and vegetables, frequently consumed yellow and orange colored soft drinks, and β-carotene-biofortified crop plants were investigated for their β-carotene and apocarotenoid contents, in a quantitative manner. The biofortified plants in this study were grains and vegetables that had been modified either through inherent genetic variability (corn (Harjes et al., 2008), sweet potato (Hotz et al., 2012), cassava (Ceballos et al., 2013)) and/or by genetic modification (rice (Paine et al., 2005; Ye et al., 2000), cassava (Welsch et al., 2010)) to contain elevated levels of β-carotene as a means to address vitamin A deficiency in at-risk populations.

The β-carotene cleavage products were present in all food items containing substantial amounts of β-carotene. The items were categorized into cereals, fruits, leafy green vegetables, non-green vegetables, and soft drinks. Cereal products represented two of the four biofortified food products examined, with rice being the most important in the Philippines. Nine replicate samples of biofortified "Golden Rice" (*Otyza sativa*) were assayed for both β-carotene and total apocarotenoid levels and found in µg/g and ng/g levels, respectively. These rice samples averaged 16.2 µg/g β-carotene and 406.1 ng/g total apocarotenoids. The biofortified corn cobs were sourced from Zambia and supplied to the investigators in unprocessed form. These five samples averaged 1.2 µg/g of β-carotene and 172.3 ng/g total apocarotenoids. Additional biofortified cassava and orange fleshed sweet potatoes from Uganda were also analyzed. The cassava samples (n=6) averaged 13.0 µg/g β-carotene and 66.2 ng/g total apocarotenoids, while the orange fleshed sweet potatoes averaged 40.9 µg/g β-carotene and 73.9 ng/g total apocarotenoids. These four biofortified sources of pro-vitamin A would represent a significant source of vitamin A activity while providing a relatively small amount of β-carotene apocarotenoids, representing, on average, ~5.0% of the β-carotene present in the biofortified rice, corn, cassava or sweet potatoes.

The locally grown and harvested (in field or greenhouse) crops in the Philippines showed similar trends in β-carotene and apocarotenoid content. Squash (n=3) and tomato (n=3) samples were found to contain 6.4 and 1.6 µg/g β-carotene, respectively, and 70.4 and 39.3 ng/g total apocarotenoids, respectively. Filipino fruits were widely tested as well, but only ripe Kinalabaw mango proved to be a significant source of both β-carotene and apocarotenoids. The three samples analyzed averaged 10.5 µg/g β-carotene and 111.2 ng/g total apocarotenoids. All of the soft drinks that were assessed showed β-carotene and apocarotenoid activity, with one brand having the highest levels of both, at 15.9 µg/g β-carotene and 103.3 ng/g total apocarotenoids.

Apocarotenoids can form through enzymatic (Harrison and Bugg, 2014) and non-enzymatic (Schaub et al., 2017) cleavage. In investigating β-carotene degradation pathways, Schaub and coworkers (Schaub et al., 2017) demonstrated significant nonenzymatic β-carotene decay over time, and they confirmed the quantitative relevance of the oxidized β-carotene polymers that the Avivagen group had previously identified (Burton et al., 2016). Using the GA assay as a marker for the presence of the β-carotene copolymers, Schaub and coworkers found that the quantity of carotenoids lost differed substantially from the quantity of apocarotenoids formed, with the latter accounting for only 0.3-0.4 percent of the initial β-carotene. This was particularly evident in the data generated from the orange fleshed sweet potatoes. Similar discrepancies were found with stored "Golden Rice" during a 215-day time course study. The β-carotene losses resulted in total amounts of apocarotenoids formed that were two orders of magnitude lower (i.e., ng *vs*. µg). The data clearly showed that, in storage, the sum of the β-carotene-derived apocarotenoids increased until ca. days 25−30, at which time ~50% of the available β-carotene was degraded. Thereafter, despite continuing β-carotene decay, the sum of the apocarotenoids remained constant, and the conclusion was reached, in accord with Burton et al. (Burton et al., 2016), that the source of the quantitative discrepancy is the natural occurrence of β-carotene copolymers. These copolymers are, therefore, the dominant product of the oxidative loss of β-carotene, and the apocarotenoids do not contribute substantially to OxBC formation, as they remain largely unchanged during the latter stages of storage.

The work of these two independent research groups underscores the fact that β-carotene copolymers are, indeed, commonly found in a variety of cereals, green and non-green vegetables and fruits, whether biofortified or not, that are consumed regularly by humans and animals.

An indication of the ubiquity and apparent safety of the β-carotene copolymers is the fact that the combined series of products that were examined by the two research groups represent products that have been used as staple foods and animal feed for centuries. Corn (maize), rice and cassava serve as the most important sources of energy for the vast majority of the world population, and corn and rice provide considerable amounts of energy for various species of livestock in six continents. This history of dietary use in human and animal diets provides further evidence of the safety of β-carotene copolymers.

**Human dietary exposure to natural sources of OxBC**

In foods containing β-carotene and other carotenoids, oxidized copolymers are present in amounts that vary widely depending upon the physical form of the source and the subsequent degree of exposure to air, heat and light during preparation (Burton et al., 2016; Schaub et al., 2017). Consequently, consumption of foods containing β-carotene and other carotenoids by humans and animals has inevitably involved a degree of exposure to carotenoid copolymer compounds over countless hundreds of years. Given that levels in some dried foods can be comparable to the original level of precursor carotenoids, the very long history of dietary exposure to these compounds provides *de facto* support for their safety at these normal levels of intake.

Nowadays, the widespread availability of dried fruit and vegetable products provides frequent and significant dietary exposure to carotenoid copolymer compounds. Common examples include copolymers present in foods containing OxBC, oxidized lycopene, and oxidized lutein. The Avivagen group described how the process of drying and powdering carotenoid-containing foods leads to the formation of carotenoid copolymers (Burton et al., 2016). Removal of moisture and cutting or grinding foods allows greater exposure to oxygen in the air, which enhances carotenoid polymer formation. Most fruits retain ~ 20% moisture when dry, while vegetables retain ~ 10% (University of Georgia Cooperative Extension Service)(University of Georgia Cooperative Extension Service). Dried foods cut into cubes or slices will contain some carotenoid polymers, while foods ground to a powder will contain more. Plant foods such as carrot, tomato, alfalfa, and sweet potato all contain high levels of carotenoids, including β-carotene, α-carotene, lycopene and lutein. These foods, and by extension carotenoid copolymers, have been part of human and animal diets for as long as carotenoid-containing fruits and vegetables have been consumed in both fresh and dried forms.

Dried vegetable ingredients are used extensively in a variety of foods by food manufacturers to prepare a variety of products. For example, carrot, tomato and sweet potato powders are often used as ingredients to prepare baby food products, including instant meals, teething biscuits and snack puffs. Several food manufacturers also provide on their websites recipes that use significant quantities of carrot, sweet potato and tomato powders as ingredients. Using the information in Table 1**,** it is possible to estimate the amounts of OxBC and total carotenoid copolymers in a variety of foods that contain significant amounts of dried vegetable or fruit ingredients.

Estimated levels of OxBC and other carotenoid copolymer products in supplements and a selection of prepared foods containing carotenoid-rich dried vegetable ingredients are provided in Tables 2 and 3. For example, baby food, baked goods, soups, stews and casseroles prepared with carrot powder can contain as much as 4-22 mg OxBC per single serving (Table 2). A recipe from Seagateproducts.com for a drink or smoothie using carrot powder is estimated to provide approximately 4-7 mg of OxBC per drink (Table 3). Foods prepared using tomato powder are especially rich sources of carotenoid copolymer compounds. For example, soups, sauces, stews and casseroles prepared with tomato powder can contain 52-78 mg of carotenoid polymer per serving (Table 2). The use of carrot, tomato and sweet potato powders is widespread and can be seen to be a source of potentially significant amounts of carotenoid copolymer products.

Several manufacturers of tomato powder describe its usefulness in preparing or substituting for tomato sauce or paste. One serving of a tomato sauce prepared from a recipe from Savory Spice Shop, for example, is estimated to provide 0.7 mg OxBC or a total of 62 mg carotenoid copolymer (Table 3). Similarly, a recipe for Hungarian goulash from Allrecipes.com that calls for tomato paste and paprika is estimated to provide at least 0.2 mg OxBC and a total of 20 mg carotenoid copolymer per serving if tomato powder is used.

It is clear from these and other examples in Table 3 that tomato powder is a rich source of carotenoid copolymers and of lycopene copolymers in particular. Lycopene being even more susceptible than ß-carotene to formation of active copolymer products (Burton et al., 2014; Johnston et al., 2014) means that the significant losses of lycopene that occur during tomato processing (Takeoka et al., 2001) will inevitably be accompanied by extensive lycopene-oxygen copolymer formation.

If foods containing tomato powder are compared (e.g., tomato sauce, soups, ketchup), there is substantially more carotenoid polymer content, 6-78 mg, arising from the large amount of lycopene originally present in the tomato, in addition to the contribution from β-carotene. As has been demonstrated elsewhere in an *in vitro* bioassay (Johnston et al., 2014), the chemically highly similar lycopene copolymer exhibits an immunological activity indistinguishable from that of the β-carotene copolymer.

Supplements are an additional source of dietary oxidized carotenoids. For example, wheatgrass powder would provide approximately 0.7 mg of OxBC and 9 mg of total carotenoid copolymer products per recommended serving (Table 2).

In ancient times, people understood the importance of preserving food for future consumption, and they had several means of accomplishing this, including drying, smoking, salting and alcoholic or acetic fermentation (Cortas, 1943). When considering the exposure to OxBC and carotenoid-oxygen copolymers, not only fresh fruits and vegetables should be considered, but preserved products especially. An expanded assessment of major sources of carotenoids, beyond that accomplished by Schaub et al. (Schaub et al., 2017) in the Philippines, reveals the further extent of these compounds in the world's food supply.

***Carrot.*** According to the United States Department of Agriculture (USDA), raw carrots contain 83 μg/g of β-carotene, 35 μg/g of α-carotene, and 88% moisture (United States Department of Agriculture, 2020). We have shown that when dried, carrots are a rich source of carotene-oxygen copolymers (Burton et al., 2016).

The close chemical identity of naturally occurring carotene copolymer, isolated from carrot powder, with the synthetic OxBC β-carotene copolymer has been established by GPC, FTIR, elemental composition and GC-MS injector port thermal decomposition analyses (Burton et al., 2016).

The earliest example of powdered carrot consumption comes from a shipwreck found off the coast of Italy (Institute for the Preservation of Medical Traditions, 2010). Built around 140-120 B.C., the ship carried medical supplies, including a tin containing a number of small green tablets. The tablets were intended to treat illnesses, and careful DNA sequencing at the *Smithsonian* revealed they contained several vegetables, including carrot.

Later, during the $12^{th}$ century, carrot powder was mentioned by the Arab writer Ibn al-Awan in his agriculture book, *Kitab al-Filahah*. He noted that carrot could make a medicinal powder when mixed with honey, date syrup and sugar. He also wrote that people of certain countries ground carrot into a powder and mixed it with flour, barley, rice or millet to make a healthy bread (Stolarczyk, 2020).

Prior to 1958 a number of patents had been filed involving dried carrot preparations as foods or ingredients. A US patent filed in 1918 described a process for producing dried carrot flakes, which were sold in packages and could be used in soups, cooked with other vegetables, or served alone after about five minutes boiling in water (Horn, 1918). Another US patent in 1928 described the preparation of vitamin supplements from dried, powdered vegetables (Prince, 1928). Carrot was one of the vegetables used, and the supplements were intended to fortify the diet with vitamins A, B, C, D and E. The vegetable powders were added to human and animal foods or compressed into tablets for subsequent sale. The invention of a dried carrot meal (i.e., powder) made from raw carrots was disclosed in a 1940 US patent (Nesbitt and Warner, 1940). This product was prepared in such a way as to retain its vitamins and minerals while also retaining good color and palatability. The patent stated that "The preservation of fruits and vegetables by drying is probably as old as civilized man, but the drying of such vegetables as carrots has never been considered satisfactory due to the loss of value and palatability in drying."

Carrot powder was described as a source of calcium in the preparation of pectin gels in a 1954 US patent (Shepherd et al., 1954), which highlighted how common a food item carrot powder was at that time. A 1957 US patent, which claimed 1951 priority in Britain, described a process for making an instant soup powder by the addition of cooked, finely divided vegetables to potato powder (Templeton, 1957). The starch in the potato powder helped to retain the vegetable flavour and aroma by preventing the loss of desired components during evaporation of the moisture. The invention was claimed to be useful for making edible food powders from fruits, meats and vegetables, such as celery, onions, beets, carrots, parsnips, swede, tomatoes, peas, cabbage, spinach, etc.

Researchers in Germany in 1907 described use of carrot soup to treat diarrhea and malnutrition in infants, as reported by Ohta et al. (Ohta et al., 1971). Their findings would become useful during World War II, when food was scarce. Japanese researchers successfully repeated the German research, as reported in papers published in 1953 and 1958, using carrot powder instead of fresh carrots to make the soup (Ohta et al., 1971). In this regard, it is interesting to note that in a trial conducted with swine, OxBC produced a significant and dose-dependent reduction in diarrhea, particularly during the early growth (starter) period (Kinh et al., 2020). It is also noteworthy that there is a dried carrot canine veterinary product marketed by Olewo, USA that is claimed to "cure dog diarrhea fast" (Olewo USA, 2017). Avivagen has analyzed the Olewo product and obtained the following results (Table 1): 315 μg/g OxBC, estimated via GA measurement; 301 μg/g isolated carotene copolymer; 228 μg/g β-carotene. These results are not dissimilar to those obtained for the orange carrot powder sample presented in Table 1. The Olewo website also claims the product "promotes healthy skin and coat', which is a benefit that has been consistently observed with Avivagen's Vivamune™ and Oximunol™ OxBC canine products.

Dried carrot has been consumed as a substitute for coffee. In the 1800s, coffee was a precious commodity and was often adulterated with various ingredients, including dried carrots, to extend it. In Germany during World War I, when genuine coffee was depleted, substitute coffee ("ersatz kaffee") helped to fill the void (Pendergrast, 2010). A number of recipes were developed with ingredients such as chicory, acorn, turnip, and carrot. "Ersatz Karotte Kaffee" was made with carrots and yellow turnips - dried roots, and it was then roasted. A 1915 US patent for a coffee substitute, which combined browned ground wheat kernels and browned ground carrot (Sattler, 1916). The mixture, when combined with water and boiled for 10-15 minutes, was claimed to produce a blend that was quite similar to real coffee.

Carrot by-products have found much utility in modern food production. The use of carrot powders is widespread in the food industry. There are two forms with distinctly different uses: i) as a food colorant and source of micronutrients, and ii) as a binder/extender. The food colorant form contains β-carotene, which imparts an orange color to food and also has until now unwittingly been providing a substantial amount of OxBC (Table 1). The binder form, known as carrot fiber when used in FDA-regulated products or as carrot isolate when used in meat and poultry products (Bolthouse Farms, 2002), is off-white in color because no β-carotene is present.

Carrot fiber is a food ingredient derived from the remnants of fresh carrots that are harvested and processed specifically for the retail carrot market. The product is an FDA-approved GRAS food additive (Bolthouse Farms, 2002) and is used in processed sauces, baked goods, bakery mixes, processed meats and other food applications. The function of the ingredient is as a binder, thickener, extender and stabilizer at use levels not exceeding 5% in the finished product.

A 2003 U.S. patent, assigned to Bolthouse Farms (Roney and Lang, 2003), describes a process to produce carrot fiber powder that was developed for the product's superior qualities in binding water and for its organoleptic properties (Roney and Lang, 2003). The product's binding feature comes from an exceptional capacity to absorb and retain water, which is useful particularly in meat products, providing for better moisture retention and keeping some items fresher longer (e.g., in breads) or increasing yield (e.g., fillers used in the meat industry). In soups and sauces, carrot fiber can be added to provide a better texture.

The invention of carrot fiber as a food additive was undertaken to fully utilize the carrot by-products of the baby peeled carrot production process. Bolthouse Farms harvests and processes approximately 1000 tons of fresh carrots on a daily basis, for the production of fresh baby peeled carrots to sell through the retail fresh produce industry. As an indication of scale, the production of carrot fiber from the by-products of the baby peeled carrot manufacturing process eliminates the disposal of approximately 100 tons of carrot waste daily (Bolthouse Farms, 2007). In a personal communication (Bolthouse-Farms, 2018), Bolthouse indicated that more than 1500 metric tons of carrot fiber powder are processed annually. Also, the carrot powder oxidizes naturally in 35 days post drying, going from orange to an off-white powder. As a result, the vitamin A level in the form of β-carotene in the stored product is less than 100 IU's/100 grams. This translates to less than 0.6 µg/g β-carotene, which is consistent with the off-white colour of the product and parallels the absence of β-carotene in the light brown carrot powder listed in Table 1.

Given that the source of the fiber is fresh carrot, it is reasonable to expect that the extensive processing and extended exposure to air will have degraded β-carotene to yield OxBC. Indeed, Avivagen has isolated carotene copolymer from a retail sample of carrot fiber powder. The level was 278 µg/g, which is similar to the OxBC β-carotene copolymer levels listed in Table 1 for orange carrot powder and the Olewo dehydrated carrot canine product.

Bolthouse Farms in their National List Petition submission to USDA for use of carrot fiber in organic foods estimated that the dietary intake of carrot fiber in a single serving of franks/sausages or meat patties/canned meat is 2.5 g/serving or 4.2 g/serving respectively (Bolthouse Farms, 2002). From our carrot powder data, this translates approximately to 1-2 mg OxBC per single item serving.

***Tomato***. Fresh tomato contains 4.5 µg/g β-carotene, 1.0 µg/g α-carotene, 26 µg/g lycopene, 1.2 µg/g lutein + zeaxanthin, and 94.5% moisture (United States Department of Agriculture, 2020). Tomato powder is a rich source of carotenoid polymer compounds. Table 1 shows that tomato powder contains more than three times the isolated carotenoid polymers than carrot powder. Although the tomato carotenoid polymers would be expected to contain β-carotene copolymers, it is expected there will be much more lycopene copolymers, which are chemically quite similar to β-carotene copolymers (Burton et al., 2016). Tomato, like carrot, is a carotenoid-rich food with a long history of consumption in dried form. Tomatoes have long been preserved in sun-dried form, starting with the Aztec empire in what is now Mexico. After the Spanish arrived in the Americas, they brought the tomato back with them to Europe, where it finds prolific use in Spanish and Italian cooking. A US patent issued in 1909 described a process for dehydrating and powdering tomatoes (Schroen, 1909). The patent detailed the advantage of dehydration over canning as a preservation technique because of decreased weight and volume, and improved preservation. Canned tomatoes needed to be eaten shortly after opening, and the amount of salicylic acid needed to preserve them exceeded the amount permitted by health authorities in most jurisdictions. Dehydrated tomato powder was sold as a powder or pressed into tablets or bars, and its taste was said to equal or exceed that of tomato puree.

Several US patents have been issued for the production of tomato soup powders. One filed in 1897 described a process for cooking and straining tomatoes and then adding flour and drying them to give an instant soup mix powder (Gere, 1901). In 1944, a patent was issued describing a process to make a tomato soup mix from tomatoes, flour or milk, beans or peas, and gelatin (Moore, 1944). The final product was sold as a powder or compressed into tablets. An improved process for making tomato soup powder was patented in 1951, where the soup mix was made by combining cooked vegetables with potato starch, as described previously for carrot soup powder (Templeton, 1957). A 1940 patent outlined how a thickened catsup could be made with dehydrated catsup or tomato powder (Nesbitt and Warner, 1940). A 1942 patent described a process for drying and exploding fruits and vegetables, including tomato (Musher, 1942), and a patent issued in 1947 proposed an improved method for making tomato powder (Derby, 1947).

Collectively, these references represent over 120 years of use of tomato products within the Western food industry and, implicitly, safe consumption by a significant fraction of the world population of the associated carotenoid copolymers.

***Sweet Potato***. Sweet potato is another staple food product that contains a large amount of β-carotene. According to USDA data, fresh sweet potato contains 85 μg/g β-carotene and 77% moisture (US Department of Agriculture, 2015). We identified the presence of OxBC in sweet potato powder (Burton et al., 2016) (Table 1) and the associated carotenoid polymers have subsequently been isolated by Schaub et al. (Schaub et al., 2017). Use of sweet potato powder, sometimes referred to as sweet potato flour, dates back to at least the mid-1800s. An 1869 US patent provided a process for peeling, cutting, drying and grinding sweet potatoes into flour (Marshall, 1869). A US patent was issued in 1884 for a sweet potato flour prepared by first baking the sweet potato, removing the skin, drying and separating the fiber, and then grinding or crushing the resultant product into flour (Whitcomb, 1884).

A US patent filed in 1916 for a coffee substitute addressed a coffee scarcity during the First World War (Brown, 1917). The substitute, made from roasted, dried ground sweet potato (95%) and cooked, dried ground velvet beans (5%), was prepared by boiling with water.

A compelling account of how common sweet potato flour was used in the US prior to 1958 is provided by G. W. Carver of the Tuskegee Institute in Alabama. In 1937, Carver published a report "How the Farmer Can Save His Sweet Potatoes and Ways of Preparing Them for the Table" (Carver, 1937). The report provided several recipes for making sweet potato flour, and listed multiple uses for the flour, including bread, cakes, pies, puddings, sauce, gravies, mock rye bread, ginger snaps, wafers, waffles, battercakes and custards.

***Pumpkin***. Raw pumpkin contains a significant quantity of carotenoids: 31 μg/g β-carotene, 40 μg/g α-carotene, and 15 μg/g lutein + zeaxanthin (US Department of Agriculture, 2015). A process for preparing pumpkin, squash and sweet potato powders, patented in 1897 (Gere, 1897), involved cooking the vegetable, adding starch to the pulp, drying, and then powdering it to produce an instant mix for making pies. The patent also mentioned that “the common article of pumpkin flour” on the market of the day, uncooked, dried pumpkin powder, required soaking and cooking prior to consumption.

***Dulse.*** The seaweed, dulse *(Palmaria palmata)*, contains significant β-carotene. According to Health Canada’s Canadian Nutrient File Database (Health Canada, 2018), raw dulse contains 31 µg/g beta-carotene and 85% moisture. Dulse is commonly eaten in Ireland, Iceland, Atlantic Canada and in the north eastern United States, either fresh or dried. Dried dulse may be eaten as is, or ground into flakes or powder. In the book, “History of Meat Alternatives (960 CE to 2014)” (Shurtleff and Aoyagi, 2014), there is a reference on pages 121-122 to a product catalog from 1938, called “The House of Better Living Catalog: Finer Natural Foods”. Based in Los Angeles, the company that published this catalog sold and shipped food products throughout the United States, including powdered dulse, dulse leaf, Irish moss, kelp, kelp-fancy, and sea lettuce.

**Livestock exposure to natural sources of OxBC**

As part of a poultry trial conducted by the Scottish Agricultural College (UK) (Pirgozliev and Offer, 2009), levels of geronic acid in a standard commercial, wheat-based poultry feed and also supplemented with 2 and 5 µg/g (ppm) of OxBC were measured by Charles River Laboratories (UK) (Ward, 2009). Levels of OxBC, estimated using the multiplication factor of 70/1000, corresponding to an assumed level of 1.4% geronic acid in OxBC, are presented in Table 4.

A natural background level of OxBC is evident in the basal diet. The mean value from six samples was $1.0 \pm 0.4$ µg/g. It is probable the naturally occurring OxBC arises from β-carotene originally present in the wheat used in the commercial diets. The average value for 2 ppm supplementation was $2.3 \pm 0.4$ ppm and for 5 ppm supplementation it was $4.6 \pm 0.7$ ppm. Given that these values for supplemented feeds also include background OxBC, the values are underestimated by about 20%, possibly reflecting the degree of uncertainty in choosing the value for the factor to convert geronic acid values into OxBC estimates.

Forage crops are rich sources of carotenoids, including, in particular, lutein, β-carotene, violaxanthin and neoxanthin (Maxin et al., 2020). Forage crop carotenoids have been reviewed by Nozière and coworkers (Nozière et al., 2006). Carotenoid levels depend upon synthesis and degradation. Synthesis occurs in plastids, mainly in leaves, which can contain 5–10 times more carotenoids than stems. Degradation occurs rapidly by oxidation, mainly through exposure to sunlight and oxygen, with substantial losses during haymaking. Moderate losses also occur during hay storage.

Grasses generally have the lowest level of β-carotene (146 mg/kg dry mass) while legumes have the highest level (438 mg/kg dry mass) (Esmail, 2020). The differences in the levels of β-carotene are mainly due to the ratio of leaf to stem in the plant and the capacity of the plant to synthesize β-carotene. Drying crops either on the ground or in barns reduces the β-carotene levels (Bruhn and Oliver, 1978). 80% of β-carotene from clover can be lost during the first 24 hours of sun-drying and becomes practically zero when the crop is dried for 4–5 days in the sun.

We now know that the losses of β-carotene that occur during drying of alfalfa are accompanied simultaneously by the appearance of corresponding quantities of OxBC (Table 1) (Burton et al., 2016).

Alfalfa has a long history as an important forage crop for animals. It is believed that alfalfa was first cultivated in ancient Iran and then introduced to Greece circa. 500 BC by invading Median armies (Griffiths, 1949; Putnam et al., 2001). Later the Romans encountered alfalfa hay when they expanded into the Eastern Caucasus Mountains and modern-day Turkey. Soon, alfalfa was grown across much of their empire because it was an ideal horse fodder to support their military.

The Romans introduced alfalfa into Europe as early as the first century AD, and during the Arab empires of the Middle Ages its use spread across Europe and North Africa (Equus Magazine, 2015). An Arabic dictionary from the 13$^{th}$ century describes alfalfa as a cultivated animal feed that is eaten fresh or dried (Wikipedia, 2020). The Spanish later brought alfalfa to the Americas during their colonization of the New World (Putnam et al., 2001). Attempts to cultivate alfalfa on the American East Coast were generally unsuccessful, due to the cold climate, high humidity and acidic soils. However, during the gold rush of 1849-1850, alfalfa seeds were imported from Chile to California, and the plant expanded rapidly across the Western United States (Putnam et al., 2001). Since the time of the gold rush, alfalfa has been an important crop in the Western half of the United States, being especially good for making hay as well as its ability to improve the quality of the soil.

After being introduced to Iowa in the late 1800's, alfalfa quickly replaced Timothy grass and clover as the preferred plant for producing hay (Living History Farms, 2021). In 1900, almost all alfalfa in the US was grown west of the Mississippi river, but the introduction of Grimm alfalfa from Germany allowed expansion of the crop 15-fold to 30 million acres in 1950 (Barr-Ag, 2012). Alfalfa was, and still is, an extremely important plant for farmers to feed to livestock. Alfalfa hay has long played a vital role in the dairy industry as an inexpensive source of nutrients for milk production. High quality alfalfa hay allowed California's dairy industry to become established, having been an important part of dairy cow rations for a long time (Feedstuffs, 2014).

Since the 1930's it was recognized alfalfa is an important source of vitamin A. Alfalfa meal was originally used for its β-carotene content and resultant vitamin A potency (Bickoff and Thompson, 1949). Dehydrated alfalfa was used in poultry rations primarily as a source of xanthophyll pigments, for example to give finished poultry a desirable yellow color (Bickoff et al., 1954), and as a source of fat soluble vitamins (carotene, vitamin E, and vitamin K) (Livingston et al., 1966).

Sun-dried alfalfa and powdered wheatgrass both contain quantities of OxBC and carotenoid copolymers, as estimated by geronic acid levels and isolation, respectively, that are a significant fraction of the original carotenoid levels (Table 1) (Burton et al., 2016). A high β-carotene content (Maxin et al., 2020) makes alfalfa a rich source of OxBC when dried into hay (Table 1), as well as being a source of other carotenoid copolymers, including lutein and neoxanthin

(Bickoff et al., 1954). It follows that other forage crop hays also will contain OxBC and carotenoid copolymer compounds.

A 1940 USDA bulletin entitled "The Uses of Alfalfa" contains information on feeding dried alfalfa to beef cattle, dairy cows, poultry, and hogs (Westover and Hosterman, 1940). It was estimated that a dairy cow consumes anywhere from 9-14 kg of alfalfa hay daily. Using the value of 61 μg/g (ppm) OxBC in alfalfa from Table 1, this would translate to 550-850 mg of OxBC daily. This can be compared to the level of 300 mg OxBC/head/d fed to dairy cattle that has recently been found to cause reductions in sub-clinical mastitis (McDougall, 2020).

Exposure to total carotenoid copolymer can be estimated using the value of 978 μg/g for carotenoid copolymer isolated from alfalfa (Table 1). A much higher value is obtained because of the presence of lutein and other carotenoids. For example, 9 kg hay x 978 μg/g would provide 8.8 g carotenoid copolymers daily.

Prior to the availability of synthetic vitamin A and xanthophylls, alfalfa, dried and ground to meal, was used for poultry and hog feeds, with 3-5% inclusion suggested for poultry and 10-15% for hogs (Griffiths, 1949). An earlier source recommended 5-7.5 % for poultry and 5-10% for hogs (Westover and Hosterman, 1940). Using the value of 61 ppm OxBC for alfalfa from Table 1 yields estimates of approximately 2-5 ppm OxBC in poultry feed and 3-9 ppm OxBC in hog feed. These estimates are close to the levels of OxBC that actually have been used successfully in feeding trials with poultry (2 ppm) (Kang et al., 2018) and swine (4-8 ppm) (Chen et al., 2020).

Estimating exposure to carotenoid copolymer using the alfalfa value of 978 μg/g (Table 1) gives values of 29-73 μg/g for poultry feed and 49-147 μg/g for swine feed.

It has been noted that "In swine feeding, dehydrated alfalfa has shown particular value in increasing breeding efficiency, improving lactation, and increasing the average number of pigs weaned per litter" (Griffiths, 1949). In a preliminary study, dietary supplementation of sows with synthetic OxBC (4-8 ppm) during the perinatal period was found to enhance the lactose concentration of sow milk, tending to increase litter weight and individual piglet weight at weaning (Chen et al., 2020). OxBC also tended to increase rates of return to estrus.

### Tables 1-4.

**Table 1.** Measured concentrations in foods of geronic acid, GA, arising from oxidation of β-carotene. GA values provide estimates of total β-carotene oxidation products, OxBC, that may be compared to levels of total carotenoid-oxygen copolymer products isolated from ethyl acetate extracts of powdered foods.

| Sample | n | Geronic Acid (GA) (ng/g) | OxBC (Calc)[a] (μg/g) | Residual β-Carotene (μg/g) (%)[b] | Isolated Carotenoid Copolymer[c] (μg/g) | Copolymer: OxBC Ratio[d] |
|---|---|---|---|---|---|---|
| Carrot juice | 4 | 12.6 ± 0.8 | 0.88 | | | |
| Carrot powder (Light brown) | 3 | 10,590 ± 550 | 741 | 0 (0%) | 756 | 1.0 |
| Carrot powder (Orange) | 3 | 5007 ± 119 | 350 | 120 (26%) | 404 | 1.2 |
| Olewo Dehydrated Carrots (Dogs) | 3 | 4500 ± 70 | 315 | 228 (42%) | 301 | 1.0 |
| Sweet potato powder drum-dried | 3 | 692 ± 22 | 48 | 45 (48%) | | |
| Sweet potato powder air-dried | 3 | 417 ± 55 | 29 | 12 (29%) | | |
| Spirulina powder | 3 | 2560 ± 10 | 179 | | | |
| Dulse seaweed powder | 3 | 1603± 39 | 112 | | 634 | 5.7 |
| Nori seaweed flakes | 3 | 2002 ±33 | 140 | | | |
| Alfalfa (sun-cured) | 2 | 869 ± 37 | 61 | | 978 | 16 |
| Wheatgrass powder | 3 | 964 ± 7 | 67 | | 924 | 14 |
| Spinach powder | 3 | 423 ± 79 | 30 | | | |
| Thyme powder | 3 | 839 ± 130 | 59 | | | |
| Tomato, raw | 5 | 1.5 ± 0.9 | 0.11 | | | |
| Tomato powder | 3 | 414 ± 46 | 29 | | 2600 | 90 |
| Tomato pomace | 3 | 113 ± 3 | 7.9 | | 1006 | 127 |
| Rosehip powder | 4 | 499 ± 12 | 35 | | 1380 | 40 |
| Cranberry, raw | 1 | 3.8 | 0.27 | | | |
| Cranberry powder | 3 | 338 ± 55 | 24 | | | |
| Paprika | 3 | 364 ± 22 | 25 | | 1080 | 42 |
| Dates, dried | 7 | 32 ± 12 | 2.2 | | | |
| Red palm oil | 3 | 60 ± 1 | 4.2 | | | |
| Milk (3.25% MF) | 2 | 6.7 ± 2.1 | 0.47 | | | |
| Milk powder (3.25% MF) | 3 | 2.0 ± 0.1 | 0.14 | | | |
| Whole egg powder | 3 | 34 ± 2 | 2.4 | | | |
| Greens Plus Original Powder | 3 | 3180 ± 132 | 223 | | | |
| GreenMin for Dogs | 3 | 3868 ± 218 | 271 | | | |
| Vega One All-In-One Nutritional Shake Powder (French Vanilla) | 3 | 137 ± 0.5 | 9.6 | | | |

[a] Estimated approximate total amount of β-carotene oxidation products, OxBC, = 70 x GA. The factor of 70 more accurately reflect GA production than the value of 50 used earlier, reflecting an estimated 1.4% geronic acid level in OxBC (Burton et al., 2016). OxBC includes mostly minor contributions from two other provitamin A carotenoids, α-carotene and cryptoxanthin. [b] Percentage β-carotene of the sum of OxBC and residual β-carotene. [c] Weight (μg) of carotenoid-oxygen copolymer fraction per gram of dehydrated food isolated by successive precipitations from ethyl acetate extract with hexane. [d] Ratio of isolated polymer fraction to OxBC. Ratio exceeds 1 because of carotenoid-oxygen copolymer compounds formed from carotenoids other than β-carotene. Tomato and rosehip are rich in lycopene; alfalfa is rich in lutein and zeaxanthin; dulse seaweed contains fucoxanthin, astaxanthin and violaxanthin

**Table 2.** Estimated levels of naturally occurring oxidized β-carotene, OxBC, and carotenoid copolymer products in some food ingredients and supplements.

| Dried Vegetable Source Ingredient | Food Uses | Vegetable Source per Serving[a] | OxBC per Serving[b] (mg) | Carotenoid Copolymer per Serving[c] (mg) | Major Carotenoid(s)[d] |
|---|---|---|---|---|---|
| Carrot Powder<br>i) orange:<br>ii) light brown: | Baby food, baked goods, soups, stews, casseroles, food colorant | 10-30 g<br>(2-6 tsp) | i) 4-11<br>ii) 7-22 | i) 4-12<br>ii) 8-22 | β-carotene, α-carotene |
| Sweet Potato Powder (drum dried) | Baby food, baked goods, soups, stews, sauces, casseroles | 8 g<br>(1 tbsp) | 0.4 | - | β-carotene |
| Tomato Powder | Soups, sauces, stews, casseroles | 20-30 g<br>(2-3 tbsp) | 0.6-0.9 | 52-78 | Lycopene, β-carotene |
| Spirulina powder | Health supplement | 7 g<br>(1 tbsp) | 1.25 | - | β-Carotene, α-carotene, echinenone, cryptoxanthin, zeaxanthin |
| Wheatgrass powder | Health supplement | 10 g<br>(2 tbsp) | 0.67 | 9.2 | Lutein, β-carotene |
| Nori seaweed flakes | Health supplement, seasoning | 3 g<br>(1 tbsp) | 0.42 | - | β-Carotene, fucoxanthin, astaxanthin, violaxanthin |
| Dulse seaweed powder | Health supplement, seasoning | 1.5 g<br>(¾ tsp) | 0.17 | 0.95 | β-Carotene, fucoxanthin, astaxanthin, violaxanthin |

[a] Per package directions. [b] Based on OxBC estimates listed in Table 1. [c] Based on isolated carotenoid-oxygen copolymer values listed in Table 1. [d] In order of relative abundance, where known.

**Table 3**. Estimated levels of OxBC and carotenoid copolymer compounds in some branded foods and food recipes.

| Branded Food or Food Recipe | Vegetable Powder (Content) | OxBC[a] (mg) | Carotenoid Copolymers[b] (mg) |
|---|---|---|---|
| Drinks and Smoothies (Seagate[c]) | Carrot (10 g per drink) | 4-7 | 4-8 |
| Warm Winter Nut Bread (Seagate[c]) | Carrot (22.5 g per loaf) | 8-17 | 9-17 |
| Organix[d] Finger Foods, Carrot Sticks (corn puffs, age 7 months +) | Carrot (2.8 g per 20 g package[f]) | 1-2 | 1-2 |
| Tomato Sauce (Savory Spice Shop[e]) | Tomato (24 g per 120 g[g]) | 0.7 | 62 |
| Sloppy Joes, Minnesota Style (The Spice House, manufacturer's recipe) | Tomato[h] (~16 g per serving) | 0.5 | 42 |
| Tomato Soup (Thrive Life) | Tomato (5 g per serving) | 0.15 | 13 |
| Ketchup (Savory Spice Shop[e]) | Tomato (2.5g per 1 tbsp) | 0.07 | 6.5 |
| Organix[d] Goodies Saucy Tomato Naughts (age 12 months +) | Tomato (1.35 g per 15 g package) | 0.04 | 3.5 |
| Hungarian Goulash (recipe: AllRecipes.com) | Paprika (1.5 g), Tomato[i] (7.1 g) per serving | 0.2 | 20 |
| Sweet Potato Pancakes (recipe: darngoodveggies.com) | Sweet Potato (~ 67 g per 3-4 pancakes[j]) | 3.2 | n.d.[k] |
| Sweet Potato Waffles (recipe: deliciousobsessions.com) | Sweet Potato (~ 33 g per 2 waffles) | 1.6 | n.d. |

[a] Amounts of OxBC per serving size estimated from OxBC levels for carrot powders, tomato powder, sweet potato powders and paprika listed in Table 1. [b]Amount per serving calculated from carotenoid copolymer levels for carrot powders, tomato powder and paprika listed in Table 1. [c] SeagateProducts.com. [d] Organix.com. [e] SavorySpiceShop.com. [f]Manufacturer's recommended serving is 5 g (3 sticks). [g] USDA serving size is 120 g or ½ cup. [h] Recipe calls for tomato powder, tomato sauce, and ketchup; amount of OxBC and copolymer calculated based on the assumption that tomato sauce and ketchup are prepared from tomato powder. [i]Recipe calls for tomato paste; amount of OxBC and copolymer calculated based on using tomato powder as a substitute, since several manufacturers recommend doing so. [j]calculated for 1/2 cup (125 mL) flour, using 8 g flour per 15 mL (1 tbsp) in Table 2. [k]not determined.

**Table 4**. Concentrations of geronic acid and OxBC in control and supplemented Starter, Grower and Finisher mash and pelleted poultry feeds determined by Charles River Laboratories (U.K.) in 30 feed samples used in trials conducted by the Scottish Agricultural College (U.K.).

| **Feed** | | **Added OxBC[a] mg/kg** | **Geronic Acid µg/kg** | **OxBC (Calc)[b] mg/kg** |
|---|---|---|---|---|
| Starter Mash | A (Control) | 0 | 14.3 | 1.00 |
| | B cornstarch | 2 | 24.7 | 1.73 |
| | C cornstarch | 5 | 60.7 | 4.25 |
| | D corncob grits | 2 | 38.2 | 2.67 |
| | E corncob grits | 5 | 71.3 | 4.99 |
| Starter Pellet | A (Control) | 0 | 23.6 | 1.65 |
| | B cornstarch | 2 | 37.7 | 2.64 |
| | C cornstarch | 5 | 65.5 | 4.59 |
| | D corncob grits | 2 | 34.8 | 2.44 |
| | E corncob grits | 5 | 76.4 | 5.35 |
| Grower Mash | A (Control) | 0 | 9.88 | 0.69 |
| | B cornstarch | 2 | 27.9 | 1.95 |
| | C cornstarch | 5 | 57.5 | 4.03 |
| | D corncob grits | 2 | 31.4 | 2.20 |
| | E corncob grits | 5 | 69.6 | 4.87 |
| Grower Pellet | A (Control) | 0 | 7.13 | 0.50 |
| | B cornstarch | 2 | 27.9 | 1.95 |
| | C cornstarch | 5 | 56.3 | 3.94 |
| | D corncob grits | 2 | 37.2 | 2.60 |
| | E corncob grits | 5 | 87.7 | 6.14 |
| Finisher Mash | A (Control) | 0 | 13.6 | 0.95 |
| | B cornstarch | 2 | 35.6 | 2.49 |
| | C cornstarch | 5 | 58.7 | 4.11 |
| | D corncob grits | 2 | 29.6 | 2.07 |
| | E corncob grits | 5 | 59 | 4.13 |
| Finisher Pellet | A (Control) | 0 | 16.5 | 1.16 |
| | B cornstarch | 2 | 24.1 | 1.69 |
| | C cornstarch | 5 | 59.7 | 4.18 |
| | D corncob grits | 2 | 41.2 | 2.88 |
| | E corncob grits | 5 | 73.2 | 5.12 |

[a] OxBC provided as a 1% premix on either a cornstarch or corncob grits carrier. [b] OxBC content calculated by multiplying geronic acid by 70, assuming a 1.4% geronic acid content in OxBC.
